# AcrVIB1 inhibits CRISPR-Cas13b immunity by promoting unproductive crRNA binding accessible to RNase attack

**DOI:** 10.1101/2024.11.17.623997

**Authors:** Katharina G. Wandera, Stefan Schmelz, Angela Migur, Anuja Kibe, Peer Lukat, Tatjana Achmedov, Neva Caliskan, Wulf Blankenfeldt, Chase L. Beisel

## Abstract

Anti-CRISPR proteins (Acrs) inhibit CRISPR-Cas immune defenses, with almost all known Acrs acting on the Cas nuclease-CRISPR (cr)RNA ribonucleoprotein (RNP) complex. Here, we show that AcrVIB1, the only known Acr against Cas13b, principally acts upstream of RNP complex formation by promoting unproductive crRNA binding followed by crRNA degradation. AcrVIB1 tightly binds to Cas13b but not to the Cas13b-crRNA complex, resulting in enhanced rather than blocked crRNA binding. However, the more tightly-bound crRNA does not undergo processing and exhibits altered target RNA binding that fails to activate collateral RNA cleavage. The bound crRNA is also accessible to RNases, leading to crRNA turnover *in vivo* even in the presence of Cas13b. Finally, cryo-EM structures revealed that AcrVIB1 binds a helical domain of Cas13b responsible for securing the crRNA, keeping the domain in an untethered state. These findings reveal an Acr that converts an effector nuclease into a crRNA sink to suppress CRISPR-Cas defense.

**Highlights:**

- AcrVIB1 binds Cas13b in the absence of a crRNA
- The bound AcrVIB1 promotes crRNA binding to Cas13b
- The crRNA binds Cas13b unproductively and is accessible to RNases
- AcrVIB1 binds the Helical-2 domain of Cas13b, preventing it from securing the crRNA

## INTRODUCTION

The ancient yet ongoing arms race between bacteria and (bacterio-)phages has led to the evolution of numerous anti-phage defense systems as well as phage-encoded counter-defenses^1–3^. One prevalent pairing comprises CRISPR-Cas systems and anti-CRISPR proteins (Acrs). CRISPR-Cas systems confer adaptive immunity by storing fragments of foreign genetic material expressed as CRISPR RNAs (crRNAs) that guide anti-phage defense by the system’s effector nuclease^4,5^. These systems possess a diverse range of molecular mechanisms that confer immune defense, with the current classification scheme dividing CRISPR-Cas systems into two classes, seven types, and over 33 subtypes and variants^6,7^. Acrs are smaller (52 - 528 amino-acids) proteins typically encoded by phages that inhibit different steps of CRISPR-based immunity^8–10^. These protein inhibitors have proven as diverse as CRISPR-Cas systems, with five CRISPR-Cas types and 13 subtypes associated with at least one inhibiting Acr^10^ as well as a wide range of inhibitory mechanisms^11–13^.

Within the diversity of inhibitory mechanisms exhibited by Acrs, the vast majority directly act on the ribonucleoprotein (RNP) effector complex formed between a Cas effector nuclease and its crRNA. In one of the most common modes across CRISPR-Cas types, the Acr binds directly to the effector complex to block target recognition or binding. As examples of this mode, AcrIF1 attaches to the hexameric Cas7f backbone within the Type I-F effector complex to block hybridization of the crRNA guide to target DNA^14,15^, AcrIIA4 occupies the protospacer-adjacent motif (PAM)-interacting site of the Type II effector nuclease Cas9 to prevent DNA interrogation^16^, and AcrVIA1 binds the guide-exposed portion of the Type VI-A effector nuclease Cas13a to block hybridization of the crRNA guide to a complementary target RNA^17^. Acrs can also cause the effector complex to non-specifically bind DNA as exemplified by AcrIF9^18^ or acetylate the effector complex or the nuclease to block recognition of the PAM flanking the crRNA target as exemplified by AcrVA5^19^. Other Acrs exert their effects on the RNP complex after the complex has bound the target. For instance, AcrIIC4 acts on the Cas9 effector complex after PAM binding but before R-loop formation^20^. Separately, AcrIIA5 acts on the Cas9 effector complex after R-loop formation by blocking the RuvC endonuclease domain^21^. Thus, a wide range of inhibitory mechanisms have been elucidated in which an Acr acts on the pre-formed RNP complex.

In contrast, only two mechanisms have emerged that solely act before RNP complex formation. In one mechanism represented by AcrIIC2 and AcrIIA17, the Acr binds to Cas9, occluding crRNA binding^22,23^. In the other mechanism represented by AcrVA2, the Acr interacts with the N-terminus of the Type V-A effector nuclease Cas12a during translation, resulting in degradation of the template mRNA^24^. While these two examples illustrate that mechanistic steps upstream of RNP effector complex formation can be inhibitory targets of Acrs, the other steps upstream of RNP formation and the many possible mechanisms-of-inhibition suggest that more paradigms await to be discovered.

In this work, we show that AcrVIB1, the only reported Acr acting against the Type VI-B effector nuclease Cas13b^25^, also acts upstream of RNP complex formation but with a unique mechanism of action: promoting rather than inhibiting binding of the crRNA to Cas13b. Binding is unproductive though, preventing processing of the precursor crRNA (pre-crRNA), reducing hybridization to a target RNA, and blocking induction of collateral RNA cleavage that normally drives cellular dormancy^26^. The bound crRNA is also susceptible to RNase attack, leading to extensive degradation in cells despite being more tightly bound by Cas13b. Finally, cryo-EM structures reveal that the Acr locks Cas13b in a flexible state accessible to partial binding of the crRNA. These findings provide molecular insights into how phages can overcome defense by Type VI-B CRISPR-Cas systems and establish a unique anti-CRISPR defense through the conversion of a Cas nuclease from an immune effector into a crRNA sink representing a dead end for the crRNA.

## RESULTS

### AcrVIB1 inhibits collateral RNA cleavage of Cas13b upstream of RNP formation *in vitro*

Our prior experiments using cell-free transcription-translation (TXTL) suggested that AcrVIB1 inhibits RNA collateral cleavage activity of PbuCas13b most strongly when acting prior to RNP complex formation^25^. To explore this observation under fully controlled conditions, we performed collateral RNA cleavage assays *in vitro* using purified components. As part of the assay (**Fig. 1A**), PbuCas13b was combined with either AcrVIB1 or an *in vitro*-transcribed crRNA. After 15 minutes, the remaining component (AcrVIB1 or crRNA) was added along with target RNA and a collateral RNA substrate linking a fluorophore and a quencher. Fluorescence stemming from separating the fluorophore and quencher through collateral RNA cleavage was then measured over time. As part of the assay, crRNAs with a processed or unprocessed terminal repeat were tested in case the processing state of the crRNAs impacts inhibition by AcrVIB1.

**Figure 1.**
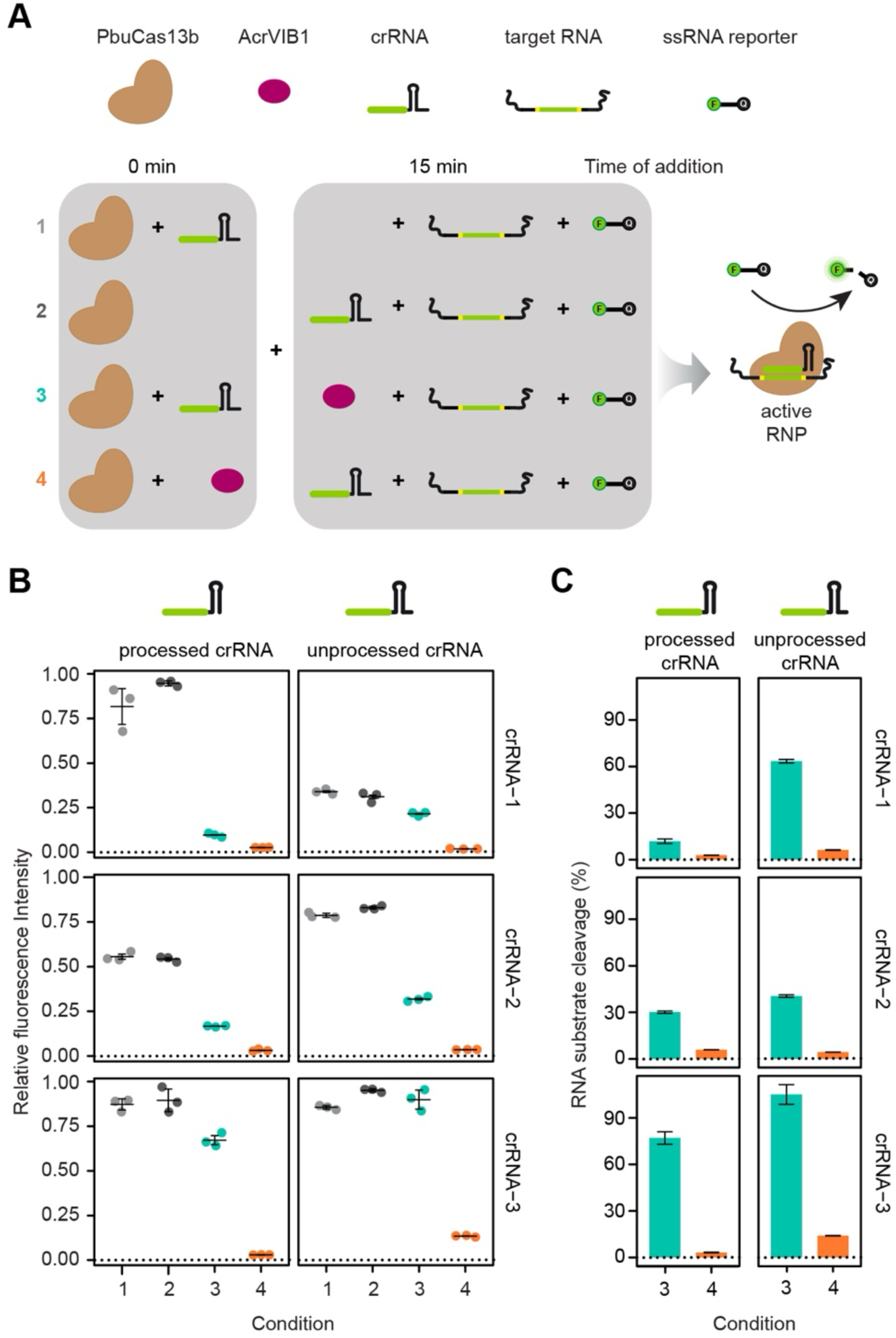
AcrVIB1 primarily inhibits RNA collateral cleavage by PbuCas13b upstream of RNP formation *in vitro*. (**A**) PbuCas13b was incubated with either a corresponding crRNA or AcrVIB1 for 15 minutes, before all other components were added. A fluorescently labeled FAM reporter and a target RNA matching the crRNA were always added last. Fluorescence was measured over time, with values shown at a selected time that ensures that the highest fluorescence signal has not saturated. See **Figure S1** for the full time courses. Incubation times are the same for a given crRNA whether processed or unprocessed (2 h for gRNA-1, 5 h for gRNA-2, and 1 h for gRNA-3). (**B**) Collateral cleavage activity of Cas13b was measured using a set of three crRNAs (crRNA-1-3). Dots represent individual fluorescence measurements, lines and error bars the mean and standard deviation of triplicate independent measurements. (**C**) Remaining collateral activity of PbuCas13b was calculated using the ratio of fluorescence measurements of conditions with and without AcrVIB1 when crRNA was either added before (3) or after (4) AcrVIB1. Values represent the ratio of the mean of fluorescence measurements and error bars represent the combined standard deviation by calculating the square root of the sum of variances.

In line with our prior TXTL results^25^, AcrVIB1 exerted a stronger inhibitory effect when added before rather than after the crRNA (**Fig. 1B**). Specifically, AcrVIB1 reduced PbuCas13b’s collateral RNA cleavage activity by 97% when added before or 88% when added after a processed crRNA-1 (**Fig. 1C**). A similar trend was observed with two other processed crRNAs, with AcrVIB1 reducing collateral RNA cleavage activity by 94% or 70% (crRNA-2) and by 97% or 23% (crRNA-3) when added before or after the crRNA (**Fig. 1B-C**). These reductions in collateral activities could not be attributed to the 15-minute difference in adding the crRNA, as collateral cleavage was similar or even slightly stronger with the delay (**Fig. 1B**). When comparing processed and unprocessed crRNAs, adding the crRNA before rather than after adding AcrVIB1 resulted in stronger inhibition for the processed versus unprocessed crRNA (12% vs. 64% for crRNA-1, 30% vs. 41% for crRNA-2, 77% vs. 105% for crRNA-3) (**Fig. 1C**), possibly due to the processed crRNA more readily binding PbuCas13b. Overall, AcrVIB1 predominantly exerts its inhibitory activity before the formation of the RNP complex.

### AcrVIB1 destabilizes the crRNA but not Cas13b *in vivo*

After confirming that AcrVIB1 principally inhibits collateral RNA cleavage by PbuCas13b upstream of RNP complex formation, we began exploring the molecular mechanism of inhibition. Prior work on the three Acrs similarly acting upstream of RNP complex formation (AcrIIC2, AcrIIA17 and AcrVA2) served as reference points, with AcrIIC2 and AcrIIA17 leading to crRNA degradation through occlusion from binding Cas9^22,23^ and AcrVA2 leading to Cas12a degradation through translational inhibition^24^. As the Acrs led to degradation of the crRNA or the Cas nuclease *in vivo*, we measured crRNA and PbuCas13b abundance when the unprocessed crRNA-1 and PbuCas13b were heterologously expressed along with AcrVIB1 in *E. coli* (**Fig. 2A**).

**Figure 2.**
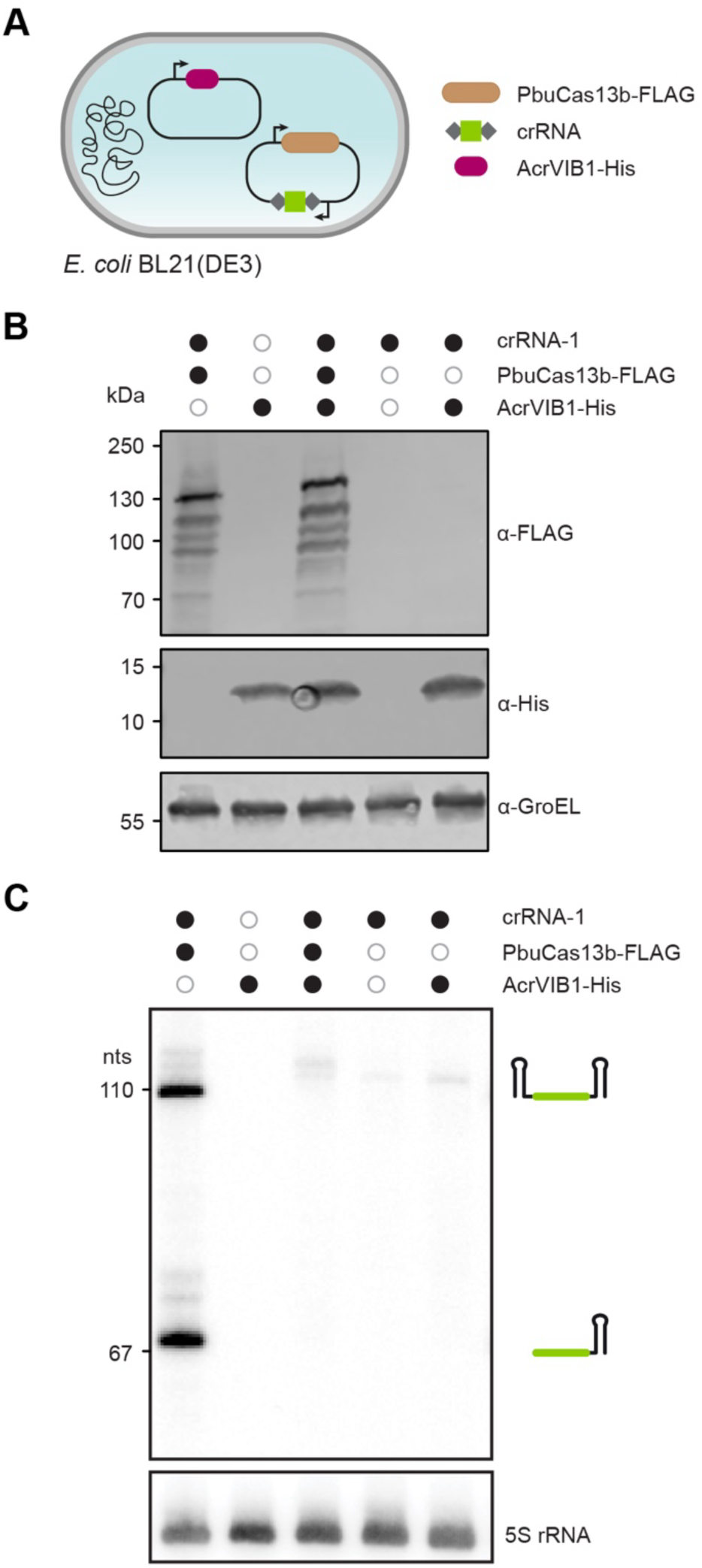
AcrVIB1 reverses crRNA stabilization by PbuCas13b *in vivo*. (**A**) Constructs expressing PbuCas13b-FLAG, AcrVIB1-His and crRNA in *E. coli* BL21(DE3). (**B**) Western blotting analysis of PbuCas13b and AcrVIB1. GroEL was detected as a loading control. (**C**) Northern blotting analysis of crRNA. 5S RNA was detected as a loading control. Gel images in B and C are representative of three biological replicates.

We first performed western blotting analysis to measure PbuCas13b levels. Co-expressing AcrVIB1 and PbuCas13b led to no difference in the expression or stability of PbuCas13b (**Fig. 2B**), ruling out a mechanism paralleling that of AcrVA2. We therefore turned to northern blotting analysis to measure crRNA levels. Here, both unprocessed and processed crRNA-1 were detectable in the presence of PbuCas13b and in the absence of AcrVIB1 (**Fig. 2C**). In contrast, both crRNA products were undetectable in the absence of PbuCas13b as well as in the presence of both PbuCas13b and AcrVIB1. Therefore, PbuCas13b protects the crRNA from turnover, which is reversed in the presence of AcrVIB1.

### AcrVIB1 binds to Cas13b and not to the crRNA

While AcrVIB1 led to destabilization of the crRNA in *E. coli*, AcrVIB1 could be interacting with either PbuCas13b or the crRNA. We therefore measured the extent of binding between AcrVIB1 and either PbuCas13b or the crRNA using microscale thermophoresis (MST) (**Figs. 3 and S3**). As part of the measurements, AcrVIB1 was tracked through its His-tag using a fluorescently tagged-Tris-NTA moiety. MST measurements showed tight binding between AcrVIB1 and PbuCas13b, with a calculated dissociation constant (K_D_) of ∼44 nM (**Fig. 3A**). In contrast, MST did not detect any binding between AcrVIB1 and the processed crRNA-1 (**Fig. 3B**). Binding between AcrVIB1 and PbuCas13b was verified by co-immunoprecipitation and mass photometry when both were incubated *in vitro* (**Figs. S2 and S4A-B**).

**Figure 3.**
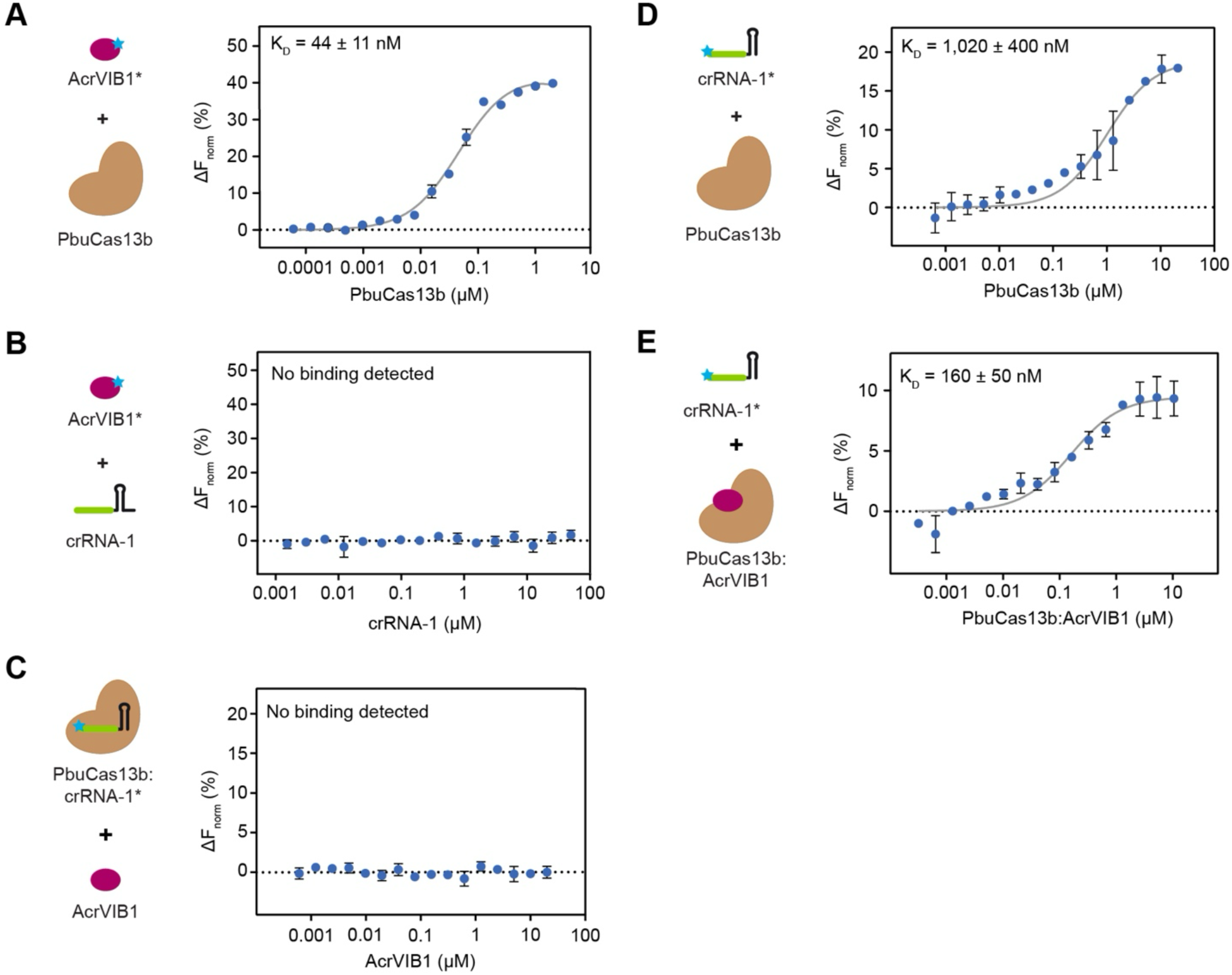
AcrVIB1 promotes crRNA binding by directly interacting with PbuCas13b. (**A-E**) Microscale thermophoresis assay to quantify interactions between (**A**) PbuCas13b and His-tag-labeled AcrVIB1, (**B**) crRNA and His-tag-labeled AcrVIB1, (**C**) AcrVlB1 and the PbuCas13b:crRNA complex, with 5′-labeled crRNA, (**D**) PbuCas13b and 5′-labeled crRNA and (**E**) the PbuCas13b:AcrVIB1 complex and 5′-labeled crRNA. In all experiments, the labeled component is indicated by an asterisk. Dots represent the movement of the molecules at a specific concentration in response to a temperature gradient. Error bars depict the standard deviation of triplicate independent measurements. The corresponding MST traces can be found in Figure S3A-E.

If AcrVIB1 tightly binds and thereby inhibits PbuCas13b while the presence of the crRNA reduces inhibition, then binding of the crRNA to PbuCas13b could negatively affect subsequent interactions with AcrVIB1. To evaluate this possibility directly, we performed MST to measure binding between AcrVIB1 and PbuCas13b already complexed with a crRNA (**Fig. 3C**). In this case, we tracked a crRNA labeled on its 5′ end with a fluorophore, which would not be removed through processing by PbuCas13b^27^. The crRNA rather than AcrVIB1 or PbuCas13b was labeled, as a binding interaction would be registered only if PbuCas13b bound both the crRNA and AcrVIB1. While no binding of AcrVIB1 was detected under this setup (**Fig. 3C**), mass photometry analysis indicated some interaction between AcrVIB1 and the PbuCas13b:crRNA complex (**Fig. S4A**). This interaction was less stable than that formed when AcrVIB1 is added first based on melting curve determination (**Fig. S4B**). Finally, an electrophoretic mobility shift assay (EMSA) showed that pre-incubating PbuCas13b with a crRNA reduces binding by labeled AcrVIB1 (**Fig. S4C-D**). Thus, we can conclude that AcrVIB1 preferentially binds to PbuCas13b rather than the PbuCas13b:crRNA complex, in line with AcrVIB1 inhibiting collateral RNA cleavage by PbuCas13b principally before RNP formation (**Fig. 1B**).

### AcrVIB1 promotes crRNA binding to PbuCas13b

After establishing that AcrVIB1 tightly binds PbuCas13b, we next asked how subsequent binding of the crRNA is affected. Following the precedent set by AcrIIC2 and AcrIIA17^22,23^, we hypothesized that AcrVIB1 occludes crRNA binding, which would explain inhibition prior to RNP formation (**Fig. 1B**) as well as AcrVIB1 yielding crRNA destabilization *in vivo* (**Fig. 2C**). To test this hypothesis, we again applied MST to measure the extent to which the labeled crRNA-1 binds PbuCas13b already complexed with AcrVIB1 (**Figs. 3A-B and S3**). Surprisingly, the crRNA bound significantly tighter in the presence rather than in the absence of AcrVIB1, with respective K_D_ values of ∼160 nM and ∼1 µM (**Fig. 3D-E**) (p = 0.032). We obtained similar results by EMSA, where crRNA binding to PbuCas13b required a high (200 nM) concentration of the nuclease and was enhanced when PbuCas13b was pre-incubated with AcrVIB1 (**Fig. S4E**). While these binding affinities were low compared to the picomolar affinities measured between guide RNAs and the *S. pyogenes* Cas9^28^, such affinities have been rarely measured and could vary widely across Cas single effector nucleases (**Fig. 1B**). Overall, in stark contrast to AcrIIC2 and AcrIIA17, AcrVIB1 enhances rather than occludes crRNA binding.

### The bound crRNA is not processed and yields altered target RNA binding

While AcrVIB1 promotes crRNA binding to PbuCas13b, the resulting complex exhibits greatly reduced target-dependent collateral RNA cleavage (**Fig. 1B**). We therefore evaluated the impact of AcrVIB1 binding on two intermediate steps between crRNA binding and collateral RNA cleavage: crRNA processing and target RNA binding. Beginning with crRNA processing, we incubated a 3′-labeled crRNA-1 with a full-length terminal repeat and downstream truncated spacer with PbuCas13b alone or already in complex with AcrVIB1 (**Fig. 4A**). While PbuCas13b alone fully processed the crRNA, PbuCas13b complexed with AcrVIB1 could no longer process the crRNA, especially at higher concentrations of AcrVIB1. The crRNA also remained intact in the presence of AcrVIB1, ruling out any degradation directly mediated by the Acr or the AcrVIB1-bound Cas13b. Thus, AcrVIB1’s interaction with Cas13b prevents crRNA processing.

**Figure 4.**
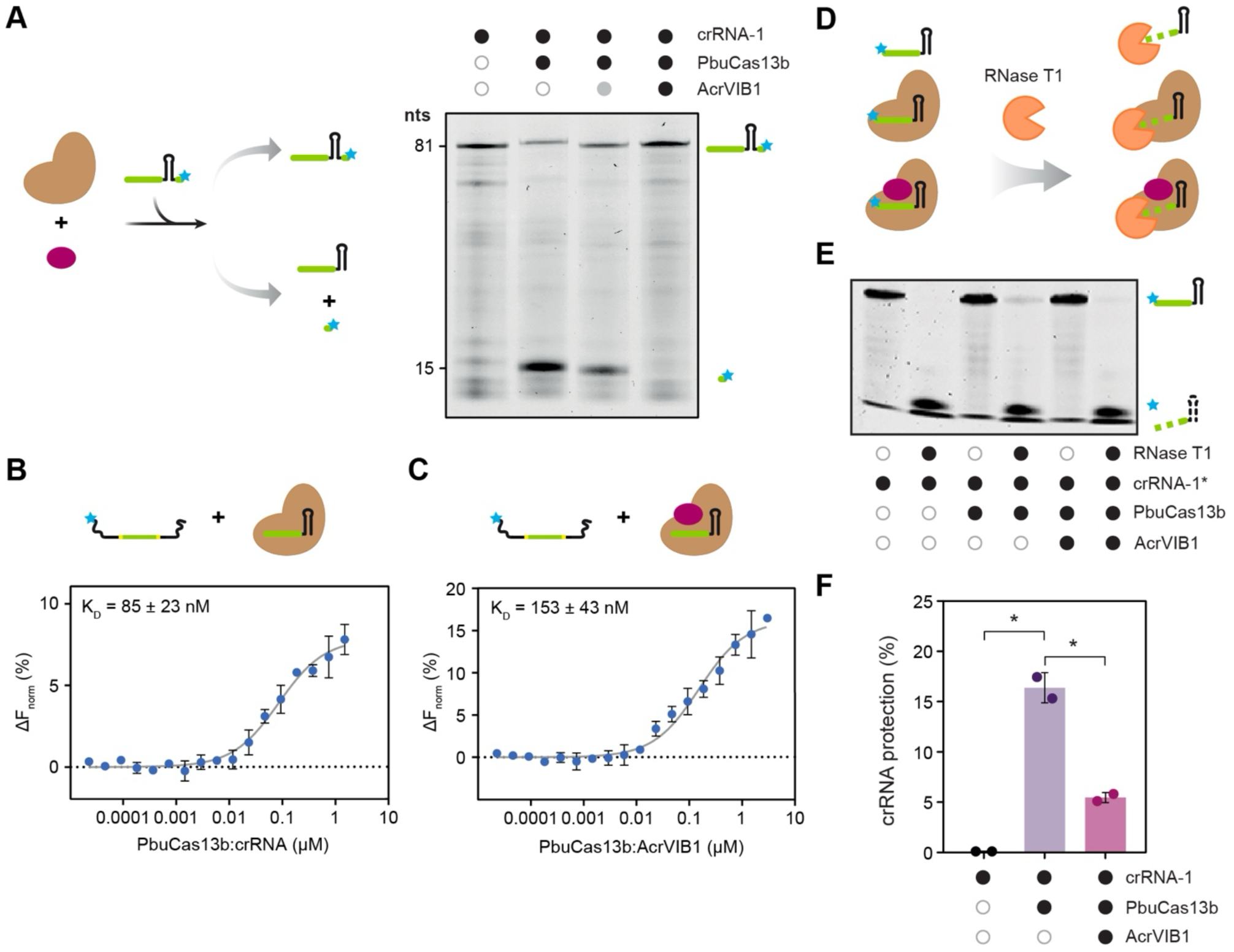
AcrVIB1 fully inhibits crRNA processing, partially inhibits target RNA binding, and facilitates crRNA degradation by an RNase even in the presence of PbuCas13b. (**A**) Assessing crRNA processing by PbuCas13b in absence or presence of AcrVIB1. AcrVIB1 was added in two concentrations indicated by the filled gray (100 nM) and black (1,000 nM) circle. The gel image is representative of two independent replicates. (**B**) Microscale thermophoresis assay to quantify interactions between the PbuCas13b-crRNA complex and a 5′-labeled target RNA. (**C**) Microscale thermophoresis assay to quantify interactions between the PbuCas13b:AcrVIB1:crRNA complex and a 5′-labeled target RNA. Buffer conditions allowed binding yet prevented cleavage of target RNA to prevent RNA turnover that could distort the measured K_D_ (see Methods). Dots represent the movement of the molecules at a specific concentration in response to a temperature gradient. Error bars depict the standard deviation of triplicate independent measurements. The MST traces corresponding to B and C can be found in Figure S3F-G. (**D**) Setup of the *in vitro* RNase degradation assay exposing PbuCas13b, AcrVIB1 and crRNA-1 to RNase T1. (**E**) PAGE analysis of 5′-labeled crRNA subjected to the RNase degradation assay. The gel image is representative of two independent replicates. (**F**) Analysis of the gel image shown in (B), using the difference of the bands between samples with and without exposure to RNase T1 to calculate the extent of crRNA protection. *: p < 0.05.

While inhibition of crRNA processing suggests disruption of other downstream steps, crRNA processing in itself is not necessary for Cas13b to recognize its target RNAs and activate collateral RNA cleavage^29^. We therefore evaluated the second intermediate step, target RNA binding (**Fig. 4B-C**). We again used MST, where we tracked the target RNA using a 5′-label. We also used a buffer that chelates free magnesium ions to allow target RNA binding without the confounding effect of collateral RNA cleavage, paralleling prior work on Cas13b^30^. Under these conditions, pre-incubating PbuCas13b with AcrVIB1 resulted in modestly but significantly reduced target RNA binding by the PbuCas13b:crRNA-1 RNP complex from a K_D_ of 85 nM to 153 nM (p = 0.047). The K_D_ value in the absence of AcrVIB1 paralleled those reported previously (27 - 42 nM)^30^. Performing a similar assay via EMSA revealed a distinct banding pattern for bound target RNA, suggesting that the interaction between PbuCas13b:crRNA and the target RNA is altered in the presence of AcrVIB1 (**Fig. S5A**). Similar changes in banding patterns for the bound target RNA as well as reduced target RNA binding in the presence of AcrVIB1 were observed via EMSA using crRNA-2 and crRNA-3 (**Fig. S5B-C**). These results indicate that AcrVIB1 affects target RNA recognition by the PbuCas13b:crRNA RNP complex, helping explain the lack of collateral RNA cleavage even in the presence of the target RNA. Between disruption of crRNA processing and altered target RNA recognition, the enhanced binding of crRNA to the PbuCas13b:AcrVIB1 can be considered unproductive.

### The presence of AcrVIB1 exposes the Cas13b-bound crRNA to RNase attack

We showed that AcrVIB1’s interaction with PbuCa13b leads to unproductive crRNA binding that prevents crRNA processing, alters target RNA binding, and stymies target-dependent collateral RNA cleavage. What remains unclear though is how these mechanisms would reduce crRNA stability *in vivo* (**Fig. 2C**), as enhanced crRNA binding to PbuCas13b in the presence of AcrVIB1 would be expected to lead to greater protection of the crRNA against host RNases. We therefore hypothesized that unproductive crRNA binding renders the bound crRNA accessible to RNases, leading to crRNA turnover *in vivo* even in the presence of Cas13b.

To test this hypothesis, we exposed crRNA-1 to RNase T1, an endoribonuclease responsible for cleaving guanine nucleotides in single-stranded RNA, in the presence or absence of PbuCas13b and AcrVIB1 (**Fig. 4D**). The crRNA alone was completely degraded by RNase T1 under our testing conditions (**Fig. 4E-F**). The presence of PbuCas13b significantly protected the crRNA from RNase T1, with ∼16% more intact crRNA than in the absence of RNase T1 (p = 0.021). We attribute incomplete protection to the poor binding affinity between PbuCas13b and the crRNA (**Fig. 3D**) that would lead to regular crRNA release. In the presence of PbuCas13b and AcrVIB1, protection significantly dropped to ∼6% compared to no RNase T1 (p = 0.021) (**Fig. 4E-F**), even though more crRNA would be bound to PbuCas13b. We therefore conclude that the crRNA is more prone to RNase degradation in the presence of AcrVIB1, explaining crRNA destabilization *in vivo* despite the Acr enhancing binding of the crRNA to PbuCas13b.

### AcrVIB1 prevents Cas13b from closing around the crRNA

To gain structural insights into the mechanism by which AcrVIB1 inhibits PbuCas13b, we determined cryo-EM structures of PbuCas13b bound to a crRNA in the absence or presence of AcrVIB1 (**Fig. 5**). When Acr was omitted, we observed a single compact structure that could be refined to 3.1 Å resolution (**Fig. 5A**) and that paralleled the previously published crystallized PubCas13b:crRNA complex (PDB entry 6DTD)^31^. Briefly, PbuCas13b adopts a heart-shaped conformation in which two HEPN domains (HEPN-1 and HEPN-2), located at the N- and C-terminus of the protein, form the apex containing the Cas13b nuclease active site, whereas two helical domains (Helical-1 and Helical-2) together with a lid-structure establish a solvent-accessible chamber that binds the 3′ stem-loop and leaves room for the unpaired 5′-stretch of the crRNA as well as for parts of the target RNA. Importantly, in this complex, the stem-loop of the crRNA is clamped tightly between the Helical-1 and Helical-2 domains, which sit on top of the HEPN-2 and HEPN-1 domains, respectively, and also contacts the lid. Consequently, the crRNA cannot leave the complex without structural rearrangements of the protein, implying that PbuCas13b adopts a more open conformation to load the crRNA. Building on prior observations garnered from the crystal structure^31^, parts of the crRNA repeat become visible in the cryo-EM structure, supporting the prior hypothesis that this stretch leaves the chamber through the large opening located at the face opposite to the active site of PbuCas13b (**Fig. 5A**).

**Figure 5.**
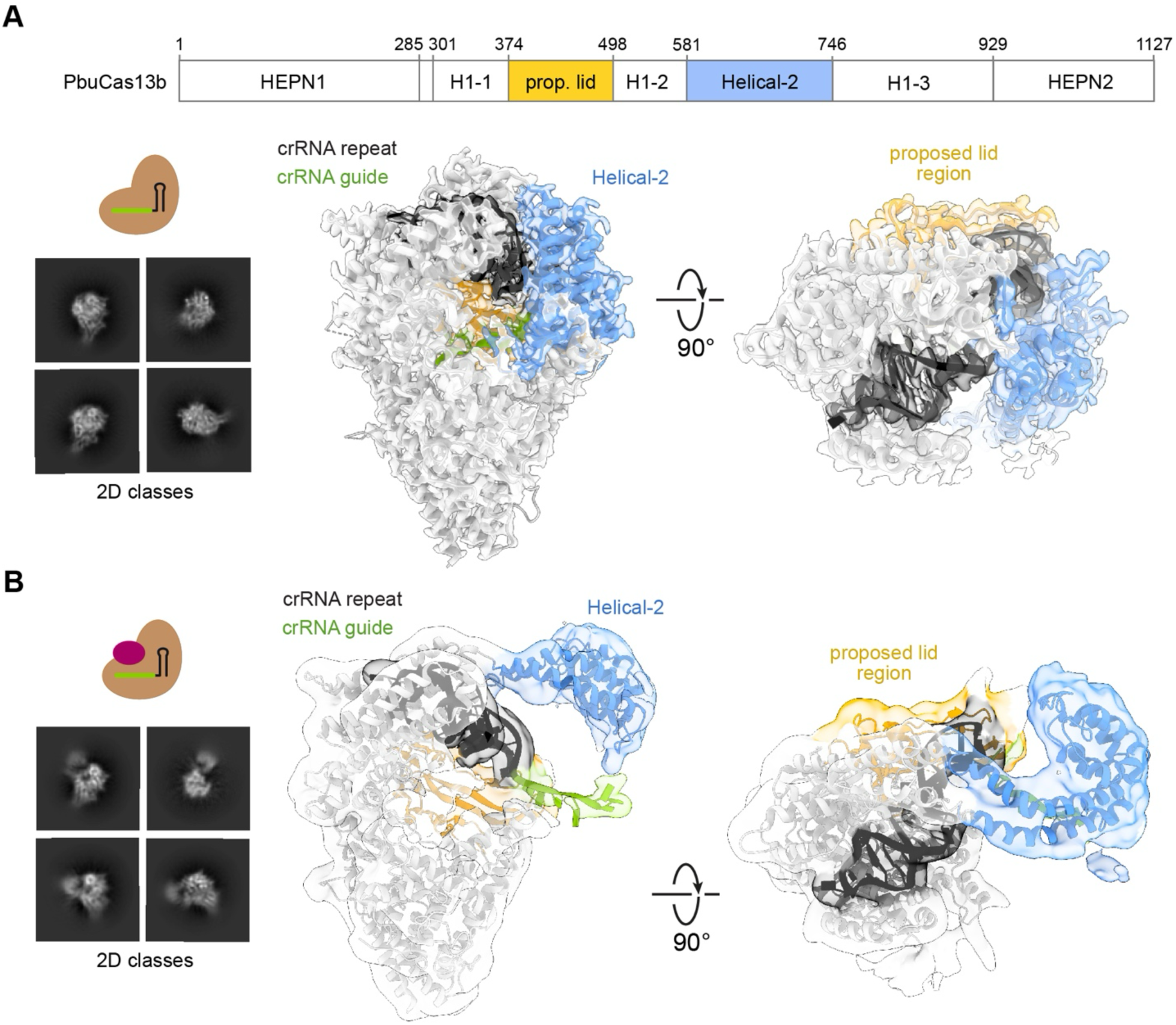
The presence of AcrVIB1 untethers the Helical-2 domain of PbuCas13b, exposing the bound crRNA. (**A**) Cryo-EM structure of PbuCas13b with bound crRNA. The structure (9FCV) has an average resolution of 3.09 Å. The cryo-EM map is colored according to the linear domain organization of PbuCas13b reported previously^31^. The crRNA is colored in green (guide) and black (repeat). Representative 2D classes of cryo-EM particles are displayed on the left. (**B**) PbuCas13b:crRNA complex in an open conformation observed in the presence of AcrVIB1. The model was obtained by morphing of the structure shown in (**A**) into a 4.23 Å resolution map. Representative 2D classes of cryo-EM particles are displayed on the left.

Including AcrVIB1 with PbuCas13b and the crRNA led to heterogeneous and less compact particles. The only trackable conformation in these samples could be refined to an overall resolution of 4.2 Å (**Fig. 5B**). While the quality of the resulting map does not reveal individual secondary structure elements, it clearly shows that the Helical-2 domain has lost its contact to HEPN-1 and the lid domains, leading to a swing-out motion of ∼60 Å at its tip. As a consequence, the crRNA is now fully solvent-exposed, and its 3′ end points away from the core of the Cas13b nuclease and into the direction of the swung-out Helical-2 domain (**Fig. 5B**).

### AcrVIB1 binds the Helical-2 domain of Cas13b that prevents the crRNA from being secured

Because this opening of the Cas13b:crRNA complex occurs in the presence of AcrVIB1, it is conceivable that the anti-CRISPR protein must be bound to the complex, thereby preventing its role as a clamp protecting and holding the crRNA in place. However, the exact position of AcrVIB1 was not obvious in the cryo-EM map because of its low resolution. While considering different possible locations, we noticed that the density around the shifted Helical-2 domain was less resolved than for the remainder of the protein. The lower resolution may be a consequence of local flexibility, capturing conformations with different degrees of opening in the map. Alternatively, the lower resolution could indicate binding of AcrVIB1 to the Helical-2 domain.

With the recent advent of AlphaFold3^32^, we explored whether AcrVIB1 interacts with the Helical-2 domain or elsewhere on PbuCas13b. As calculations with the complete nuclease and AcrVIB1 were inconclusive, we modeled a complex of AcrVIB1 only with the Helical-2 domain. In contrast to similar approaches with the predecessor software AlphaFold2^33^, the new version resulted in a reproducible high-confidence model of a 1:1 complex (**Fig. 6A**). When this model is mapped to the open conformation of the Cas13b:crRNA complex, AcrVIB1 wedges between the lid and the Helical-2 domain (**Fig. 6F**). Importantly, when the Helical-2:AcrVIB1 model is superimposed onto the closed conformation of PbuCas13b, AcrVIB1 would arrive at the inner RNA-binding cavity and lead to severe clashes with parts of the nuclease, explaining why AcrVIB1 does not bind to the pre-formed PbuCas13b:crRNA complex in our experiments. On the other hand, the open conformation is expected to allow the crRNA to more easily access its binding site in PbuCas13b, which could explain why AcrVIB1 strengthens crRNA binding. At the same time, the crRNA remains unprotected in this complex, making it prone to degradation by RNases as we observed experimentally (**Fig. 4D-F**). The inability of the Helicase-II domain to close around the crRNA likely prevents further conformational changes previously hypothesized as necessary steps to activate collateral RNA cleavage^31^.

**Figure 6.**
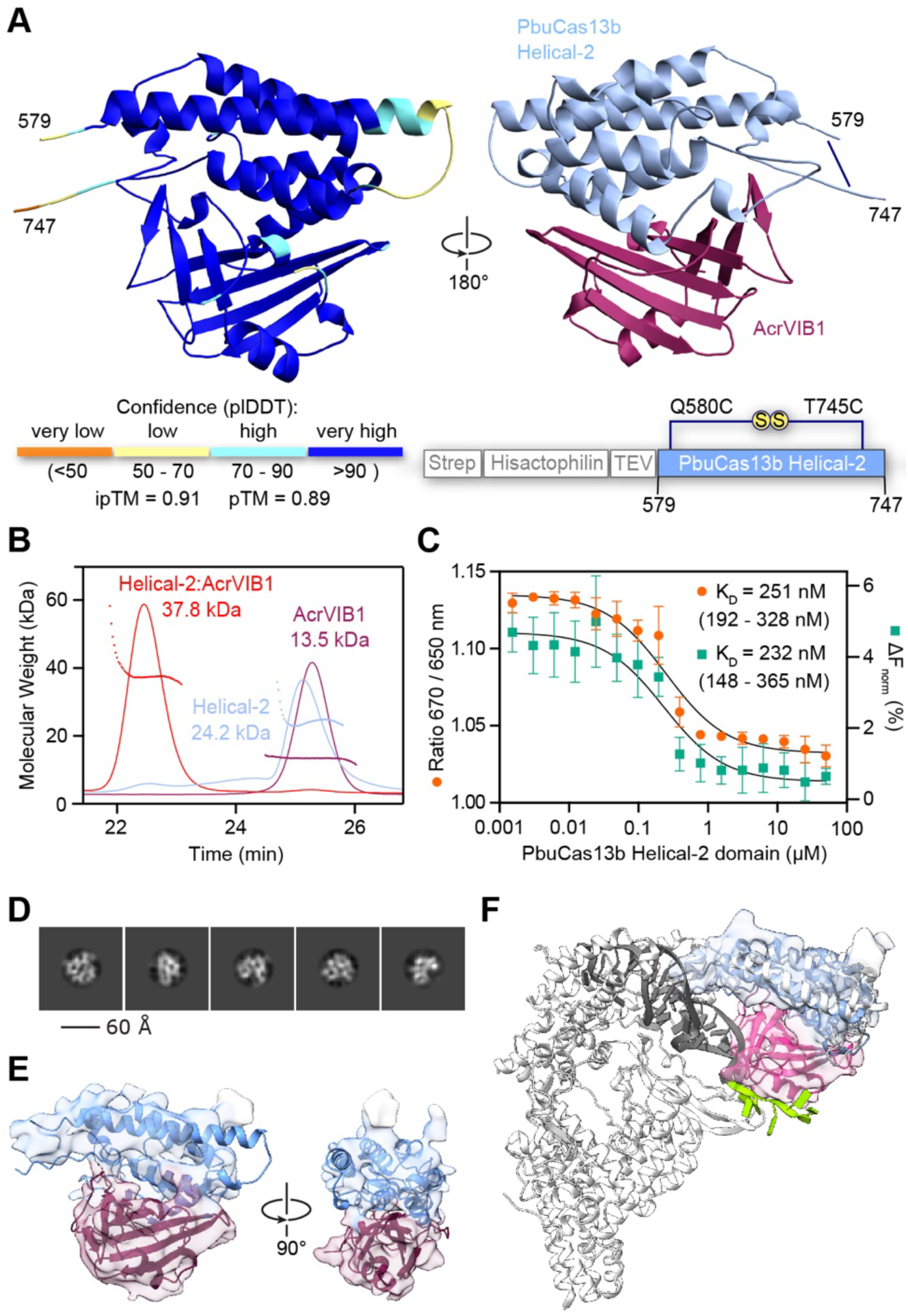
AcrVIB1 binds the Helical-2 domain of PbuCas13b. (**A**) AlphaFold3 model of the PbuCas13b Helical-2 domain (residues 579 - 747) bound to AcrVIB1. Left: structure colored according to the modeling confidence (plDDT score). Right: 180° rotation of the model and colored to indicate AcrVIB1 (violet) and the Helical-2 domain (blue). A dark blue line indicates the position of an artificial disulfide bond that was introduced for stabilization of the isolated Helical-2 domain. (**B**) SEC-MALS profiles of AcrVIB1 (violet), the PbuCas13b Helical-2 domain (blue) and the complex of both (red). The obtained molecular weights are in good agreement with the expected values (AcrVIB1: 13.6 kDa, PbuCas13b Helical-2 domain: 20.4 kDa, complex: 34.0 kDa). Results are representative of duplicate independent runs. (**C**) Binding-affinity measurements of the PbuCas13b Helical-2 domain to fluorescently labeled AcrVIB1. Orange dots: analysis based on the 670 / 650 nm emission ratio of the fluorescent dye. Teal squares: analysis of the same samples based on MST. Dots/squares and error bars represent the mean and S.D. of 4 independent replicates. (**D**) Representative 2D classes of the 34-kDa complex comprising PbuCas13b Helical-2 domain (residues 579 - 747) and AcrVIB1. (**E**) Superposition of the AlphaFold3 model from (**A**) with the 34-kDa AcrVIB1:PbuCas13b Helical-2 domain (residues 579 - 747) complex. The resulting map has a resolution of 5.11 Å. (**F**) Full-length PbuCas13b (light gray) with bound cRNA (repeat in dark gray, guide in green) and an open and extended Helical-2 domain superposed with the cryo-EM complex shown in (**E**). AcrVIB1 is colored in violet and PbuCas13b Helical-2 domain is colored in blue.

To validate the AlphaFold3 model experimentally, we produced the Helical-2 domain recombinantly and assessed complex formation with AcrVIB1 by SEC-MALS and Spectral Shift titration. This analysis revealed a 1:1 complex of the expected molecular mass (**Fig. 6B**), with a dissociation constant of 251 nM (**Fig. 6C**). While the recombinant complex was recalcitrant to crystallization, we could corroborate the overall shape and position of the predicted secondary structure elements with a low-resolution cryo-EM reconstruction of the 34-kDa Helical-2:AcrVIB1 particle (**Fig. 6D-E**).

To probe the importance of residues in the predicted interface between PbuCas13b and AcrVIB1 (**Fig. S6A**), we tested associated amino-acid substitutions in both PbuCas13b and AcrVIB1 using our previously reported TXTL assay^25^. The assay reads out collateral activity through silencing of a targeted GFP transcript, which is reversed through inhibition by the expressed AcrVIB1 (**Fig. S6B**). Of the three tested substitutions in PbuCas13b (K627A, N634A, R662A) and AcrVIB1 (D37A/D38A, S56A, T70A), R662A in PbuCas13b as well as D37A/D38A and T70A in AcrVIB1 mediated a substantial reversal of inhibition (**Fig. S6C-D**). The attenuated inhibition supports the predicted interactions between AcrVIB1 and the Helical-2 domain of PbuCas13b, while the lack of inhibition at the other tested residues could reflect multisite interactions that collectively contribute to binding and inhibition.

As an additional means to probe binding interactions, we evaluated the *Riemerella anatipestifer* (Ran)Cas13b, which shares 43.4% identity with PbuCas13b. RanCas13b and AcrVIB1 coexist in the same *R. anatipestifer* genome, although AcrVIB1 poorly inhibits this Cas13b^25^. We hypothesized that the few amino-acid differences within the implicated portion of the Helical-2 domain prevents inhibition by AcrVIB1 (**Fig. S6E**). Substituting four residues in RanCas13b to match the aligned residue in PbuCas13b (K559N, Y609K, W627R, R630Q), we found that one substitution (R630Q) substantially enhanced inhibition by AcrVIB1 in the TXTL assay (**Fig. S6F**). Sequence divergences in this region of the Helical-2 domain thus appear to limit the host range of AcrVIB1. Overall, our mutational analyses help corroborate AcrVIB1 directly binding to the Helical-2 domain of PbuCas13b.

## DISCUSSION

Based on our collective results, we propose a detailed model governing the mechanism of action of AcrVIB1, the only known inhibitor of a Cas13b nuclease (**Fig. 7**). In the absence of the Acr, Cas13b binds and processes the pre-crRNA, after which target RNA can be recognized to activate collateral RNA cleavage. In the presence of the Acr, AcrVIB1 tightly binds Cas13b through the Helical-2 domain, thereby locking Cas13b in an open conformation that more readily allows binding of a pre-crRNA. pre-crRNA binding is unproductive though, preventing pre-crRNA processing and leading to altered target RNA binding that does not activate collateral RNA cleavage even in the presence of the target RNA. Additionally, the crRNA bound to the Cas13b:AcrVIB1 complex is accessible to RNases, making it susceptible to at least partial degradation. If the degraded crRNA is released, the Cas13b:AcrVIB1 complex could subsequently bind but not protect additional crRNA molecules, effectively turning Cas13b into a crRNA sink. Alternatively, if the untouched portion of the crRNA remains bound to the Cas13b:AcrVIB1 complex, then release of AcrVIB1 could lead to formation of a closed Cas13b:crRNA complex, but without the guide region necessary for target recognition. In either case, AcrVIB1 blocks immune activation of Cas13b in the presence of a target RNA. If a crRNA is already bound to Cas13b, then AcrVIB1 loses its ability to inhibit immune activation because the binding surface on the Helical-2 domain is buried in the Cas13b:crRNA complex.

**Figure 7.**
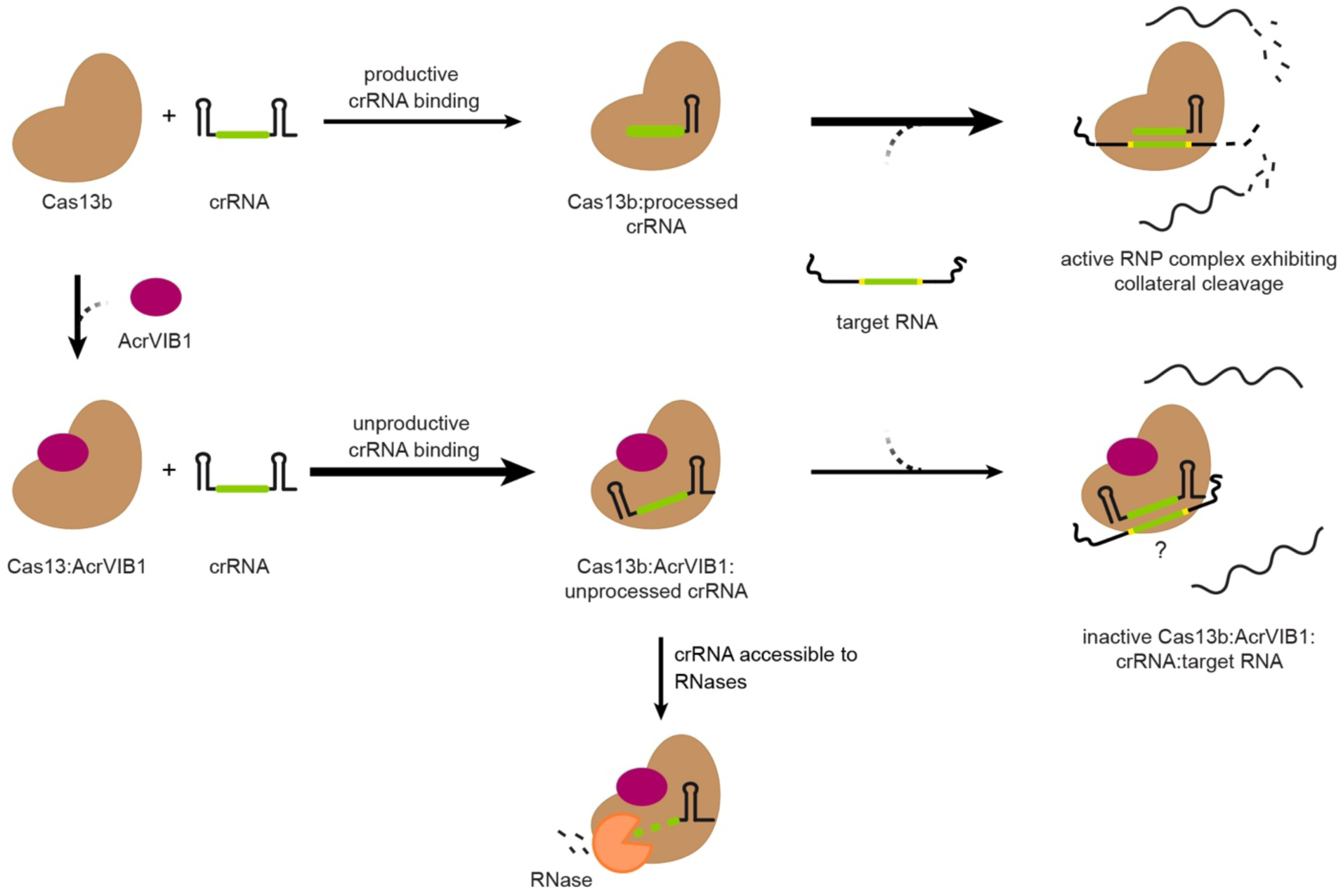
Proposed model for inhibition of Cas13b-based immunity by AcrVIB1. Cas13b weakly binds to a crRNA to form an RNP complex followed by crRNA processing. The RNP complex can then recognize a target RNA that activates collateral RNA cleavage. AcrVIB1 binds to Cas13b, leading to stronger crRNA binding to the Cas13b:AcrVIB1 complex. The bound crRNA does not undergo processing and yields altered target RNA binding that prevents activation of target-dependent collateral RNA cleavage. The crRNA bound to the Cas13b is protected from RNases, while the crRNA bound to the Cas13b:AcrVIB1 complex is susceptible to RNase attack.

Apart from supporting the proposed inhibitory mechanism of AcrVIB1, the cryo-EM analyses shed light on the crRNA loading step of Cas13b. This step has previously remained obscure due to the fact that only two crystal structures of Cas13b:crRNA complexes are currently available, and these structures do not provide direct evidence for interdomain flexibility within Cas13b nucleases^31,34^. Binding of AcrVIB1 revealed an alternative conformation in which the Helical-2 domain remains in a swung-out conformation. As this domain directly interacts with the crRNA in the Cas13b:crRNA structures, we propose that it opens to accommodate crRNA binding. It is conceivable that the same domain also moves to accommodate complete hybridization between the crRNA and target RNA, although the previously proposed lid domain could also serve this function^31^. Thus, there are still many features of Cas13b activation that remain unclear, such as how target recognition leads to nuclease activation. Further studies with AcrVIB1 could help dissect these steps.

The elucidated mechanism of action represents a distinct entry into the short but diverse list of known Acrs acting upstream of RNP formation. Among already known mechanisms are the degradation of effector protein mRNA and the occlusion of crRNAs to prevent RNP complex formation^22–24^. Separate from Acrs, RNA-based anti-CRISPR elements called Racrs were recently shown to act upstream of RNP formation by mimicking crRNA repeats that form non-functional RNPs^35^. AcrVIB1 introduces a distinct mechanism to this list by promoting rather than occluding crRNA binding, but in a way that blocks target-dependent activation and leads to crRNA degradation *in vivo*. At most, this mechanism resembles that of AcrVA1, which cleaves the guide region of a Cas12a crRNA^36^. However, AcrVA1 only cleaves the crRNA within the formed RNP, while AcrVIB1 promotes the formation of a non-functional RNP. Given the diverse mechanisms exhibited by AcrVIB1 and these few other Acrs and Racrs, additional Acrs likely await discovery. For instance, these hypothetical Acrs could act through similar mechanisms against other Cas nucleases besides Cas13b or act at other steps upstream of RNP formation, such as transcription of the Cas nuclease or pre-crRNA, crRNA processing by host factors such as RNase III or RNase E, or as part of spacer acquisition.

Given that AcrVIB1 and these other examples would not be effective after the RNP effector complex is formed, it raises the question: under which infection conditions do these examples help circumvent CRISPR-based immunity? Inhibition would be unlikely to benefit an initial infection by an Acr-encoding phage, as the expressed Acr (or Racr) would not inhibit pre-formed RNP complexes surveilling for the invading phage genome. Instead, these Acrs could assist with a follow-on wave of phage infections, an altruistic effect also associated with Acrs inhibiting some DNA-targeting CRISPR-Cas systems^37,38^. If the phage contains a mutated target site but the site could drive priming^39,40^, then Acrs acting upstream of RNP formation could inhibit CRISPR immunity before a new spacer can be acquired and produce a matching crRNA. As Acrs are often found together in anti-defense islands, multiple mechanisms of inhibition acting upstream of and on RNP complex formation could also yield more robust anti-defense under different infection conditions. Overall, more work exploring the ramifications of different inhibitory mechanisms of Acrs under different infection conditions could shed new light on the bacterial-phage arms race.

One notable observation while elucidating AcrVIB1’s mechanism of action is the poorer binding affinity between PbuCa13b and its crRNA, at least compared to that measured for SpyCas9^28^. We still observe collateral RNA cleavage activity, so the low binding affinity between PbuCas13b and crRNA does not compromise RNP activation. This observation raises the question of how variable binding affinities can be between Cas effector nucleases and their crRNAs and their potential consequences on CRISPR immunity. In the case of Cas13b and AcrVIB1, we speculate that the poor binding affinity grants AcrVIB1 an opportunity to inhibit Cas13b before a crRNA is bound; otherwise, AcrVIB1 would be rendered ineffective both to bind Cas13b and serve as a crRNA sink.

Beyond its relevance to bacterial-phage interactions, the identification and characterization of AcrVIB1 lays the foundation for its use for controlling Cas13-based functions. Indeed, Cas13 nucleases have been adapted for a number of RNA-centric applications, including post-transcriptional silencing, RNA editing, molecular diagnostics, and *in vivo* imaging^41–44^. Currently, there are only a limited number of Acrs that inhibit Type VI CRISPR-Cas systems, and only one published application using the Acr for phage engineering^45^. In this context, the Cas13a inhibitor AcrVIA1 is introduced as a selection marker via homologous recombination, together with a gene-of-interest^17^. Only phages carrying the gene-of-interest and AcrVIA1 would survive Cas13a targeting due to the inhibitory activity of AcrVIA1 against Cas13a^45^. As AcrVIB1 acts prior to RNP complex formation, applications that rely on a pre-formed RNP complex would be less suitable, such as phage engineering or molecular diagnostics. However, there are other applications in which an RNP complex is not pre-formed. For instance, as part of post-transcriptional silencing, AcrVIB1 could be applied to reduce leaky expression prior to Cas13 induction^43,46,47^, or it could restrict silencing to specific cell types through tissue-specific expression of the Acr^48,49^. Thus, further exploring the potential uses of Cas13 Acrs could expand and enhance current capabilities of CRISPR technologies.

## ACKNOWLEDGMENTS

We express our gratitude to the protein expression facility of the Rudolf-Virchow-Zentrum (Center for Integrative and Translational Bioimaging) for their excellent support and expertise in protein purification. We would also like to thank Ute Widow for excellent technical assistance in protein production. Our appreciation also goes to Thomas Gaudin for assistance with experiments using the Echo 525 acoustic liquid handler and to Mitchell O’Connell for initial discussions and technical support.

## AUTHOR CONTRIBUTIONS

Conceptualization: K.G.W., C.L.B.; Experiments: K.G.W., S.S., A.M., A.K., P.L., T.A.; Writing - Original Draft: K.G.W., S.S., P.L., W.B., C.L.B., Writing - Review & Editing: all authors; Visualization: K.G.W., S.S., P.L., W.B., C.L.B.; Supervision: N.C., W.B. and C.L.B.; Funding acquisition: N.C., W.B., C.L.B.

## MATERIALS & METHODS

### Buffers, plasmids and oligos

All resources are listed in **Table S2**.

### Molecular cloning

All plasmids were cloned using standard cloning techniques: Q5 mutagenesis, Gibson Assembly and Golden Gate Cloning. Cloned sequences were verified by Sanger sequencing or Nanopore sequencing. Descriptions and links for each plasmid can be found in **Table S2**.

### Protein purification

The gene encoding for AcrVIB1 was cloned into a modified pCOLA Duet-1 vector (Novagen) encoding for an N-terminal Hisactophilin tag^50^ and TEV-protease recognition site (KW-426). The gene encoding for PbuCas13b was cloned into pETM11 together with N-terminal hexahistidine and SUMO tags (KW-241). A construct for the Helical-2 domain of PbuCas13b was designed comprising the residues 579 - 747 and the mutations Q580C and T745C in order to form an artificial disulfide bridge for stabilization of the isolated domain. The construct was sourced commercially as a synthetic gene (GenScript) together with a Strep-tag II, a Hisactophilin tag^50^ and a TEV-protease recognition site at the N-terminus and cloned into pET28a (PL-001).

All proteins were expressed in *E. coli* BL21 (DE3) Star in baffled shaking flasks with 1 L of ZYM-5052 auto-inducing medium^51^ each at 20°C for 20-24 h.

For the purification of AcrVIB1, the cell pellet was resuspended in a buffer containing 20 mM HEPES/NaOH pH 7.5, 300 mM NaCl, 1 mM TCEP, 20 mM imidazole and one tablet of cOmplete EDTA-free protease inhibitor cocktail (Roche). After lysis by sonication, the protein was isolated from the supernatant after centrifugation for 1 h at 100,000 x g using a 5 ml HisTrap HP column (Cytiva) and eluted using a linear gradient towards 500 mM imidazole. The affinity tag was cleaved off with TEV protease (1:50 mg/mg) at 4°C during overnight dialysis against imidazole-free buffer. Gel filtration was carried out using a HiLoad 16/600 Superdex 200 pg column (Cytiva) in 20 mM HEPES/NaOH pH 7.5, 300 mM NaCl, 1 mM TCEP. The peak fractions were concentrated, flash-frozen in liquid nitrogen and stored at -80°C.

For the preparation of PbuCas13 for cryo-EM analysis, the cell pellet was resuspended in a buffer containing 50 mM HEPES/NaOH pH 7.5, 500 mM NaCl, 0.5 mM DTT and 20 mM imidazole, plus one tablet of cOmplete EDTA-free protease inhibitor cocktail (Roche). After lysis by sonication, the protein was isolated from the supernatant after centrifugation for 1 h at 100,000 x g using a 5-ml HisTrap HP column (Cytiva) and eluted using a linear gradient towards 250 mM imidazole. The affinity and SUMO tags were cleaved off with SENP2 (1:50 mg/mg) at 4°C during overnight dialysis against SEC buffer (20 mM HEPES/NaOH pH 7.5, 300 mM NaCl, 0.5 mM DTT). Gel filtration was carried out using a HiLoad 16/600 Superdex 200 pg column (Cytiva) in 20 mM HEPES/NaOH pH 7.5, 300 mM NaCl, 1 mM TCEP. The peak fractions were concentrated and subjected to cation exchange chromatography using a 1-mL HiTrap Capto SP ImpRes column (Cytiva) with a buffer containing 20 mM HEPES/NaOH pH 7.0, 400 mM NaCl and 2 mM DTT. The protein was eluted using a linear gradient from 400 mM NaCl to 1 mM NaCl. Peak fractions were pooled, resulting in a final NaCl concentration of 620 mM in the buffer. The samples were flash-frozen in liquid N_2_ for cryo-EM analysis. For cryo-EM analysis, PbuCas13 samples were diluted to 1 mg/ml in 20 mM HEPES/NAOH pH 7.0, 120 mM NaCl prior complex formation.

For biophysical analyses of PbuCas13b, cell pellets were resuspended in lysis buffer (20 mM HEPES/NaOH pH 7.5, 500 mM NaCl, 10 mM Na_3_-citrate, 10 mM MgSO_4_ and 5 mM DTT) prior to sonication. Affinity purification was conducted using several steps of lysis buffer with increasing imidazole concentrations. The desired protein eluted between 62.5 and 100 mM imidazole. Fractions of 2.5 ml were collected into tubes pre-filled with 5 ml of imidazole- free buffer. After cleavage with SENP2, anion exchange was performed using a 5 ml HiTrap Heparin column (Cytiva) in lysis buffer with a linear gradient from 0.5 M – 2 M NaCl. The elution peak was concentrated and subjected to gel filtration using a HiLoad 16/600 Superdex 200 pg column (Cytiva) in 20 mM HEPES/NaOH pH 7.5, 300 mM NaCl and 5 mM DTT. Peak fractions were concentrated and flash-frozen in liquid nitrogen.

For the purification of the PbuCas13b Helical-2 domain, the cell pellet was resuspended in a buffer containing 20 mM HEPES/NaOH pH 7.5, 150 mM NaCl, 20 mM imidazole and one tablet of cOmplete EDTA-free protease inhibitor cocktail (Roche). After lysis by sonication, the protein was isolated from the supernatant after centrifugation for 1 h at 37,000 x g using a 5 ml HisTrap HP column (Cytiva) and eluted using steps of increasing imidazole concentrations. The protein eluted at a concentration of 500 mM imidazole. The affinity tag was cleaved off with TEV protease (1:50 mg/mg) at 4°C. Gel filtration was carried out using a HiLoad 16/600 Superdex 75 pg column (Cytiva) in 20 mM HEPES/NaOH pH 7.5, 150 mM NaCl. The peak fractions were concentrated, flash-frozen in liquid nitrogen and stored at -80°C.

All chromatography steps were executed using an Äkta Go chromatography system (Cytiva).

### RNA isolation

Total RNA was isolated using a phenol-chloroform extraction method. Bacterial cultures were cultivated until reaching a maximum optical density (OD) of 1, and subsequently harvested to obtain an overall OD of 4. This was achieved, for instance, by collecting 4 mL of culture with an optical density of 1. A mixture of 95% EtOH (Sigma-Aldrich, 32221-M) and 5% phenol (Carl Roth, #A980.1) was added to each culture in a volume ratio of 1:5 from the initial culture volume, which was then immediately transferred to liquid nitrogen. Samples were placed on ice until all samples were thawed (45-60 min). Cells were pelleted at 7,400 rpm for 20 min at 4°C. Supernatant was carefully removed before centrifuging all samples again at 7,400 rpm for 1 min at 4°C. All remaining supernatant was removed. All samples were resuspended using 600 µL of 0.5 mg/ml lysozyme (Carl Roth, #8259.2) solved in TE buffer pH 8.0. After transferring all samples to a 2 mL tube, 60 µL of 10% w/v SDS was added and mixed by inversion. Tubes were placed in a water bath at 64°C for 1-2 min (the samples should become clear). To each sample, 66 µL of 3 M NaOAc, pH 5.2 was added and mixed by inversion. Equal volumes (750 µL) of phenol (Carl Roth, #A980.1) were added to the samples, and the samples were incubated for 6 min at 64°C. Samples were inverted every 30 s during the incubation process. Subsequently, all tubes were chilled on ice, before they were centrifuged at full speed (13,000 rpm) for 15 min at 12°C. The aqueous layer was transferred into 5PRIME Phase Lock Gel (PLG) tubes (VWR International, #733-2478). An equal volume (750 µL) of chloroform (Roth, #Y015.2) was added in each PLG tube and mixed by inversion. All samples were spun down at full speed for 15 min at 12°C. After that, the aqueous layer was transferred into 2 mL of RNAse-free tubes. To each sample, 2.5 volumes of a 30:1 mix of EtOH:NaAcetate (3 M, pH 6.5) was added. Samples were incubated at -80°C for at least 20 min up to 2 h. Subsequently, all samples were spun down at 15,000 g for 30 min at 4°C. The extracted RNA was washed with 500 µL of 70% EtOH and spun for 10 min at 15,000 g at 4°C. The wash step is repeated using 100% EtOH. All EtOH was aspirated and the RNA samples were dried at room temperature for 20 min before resuspending the RNA in 50 to 100 µL of RNase-free water. The concentration of each extracted RNA was determined using a Nanodrop.

### *In vitro* cleavage assays

Collateral RNA degradation is measured using a fluorescence-based assay, where a labeled RNA FAM reporter is added together with purified PbuCas13b, a corresponding crRNA and the matching target RNA. Different combinations of components were added together and incubated at room temperature for 15 min. All remaining components were subsequently added. The FAM reporter was always added last. All shown data was produced using the Echo525 Liquid Handling system. A plate reader was used to measure fluorescence over time. Raw data was imported into R (R version 4.2.0 (2022-04-22)) from .csv files using the readxl package (version readxl_1.4.3). Data wrangling was performed via the R packages dplyr (version dplyr_1.1.4), tidyr (version tidyr_1.3.0), and stringr (version stringr_1.5.1). Data of multiple experiments was merged and fluorescence intensities for each experiment were scaled to values between zero and one (minmax). Endpoints were selected manually to show data below maximum fluorescence intensity. Data visualizations were created using ggplot2 (version ggplot2_3.4.4). Calculation of the remaining collateral activity of PbuCas13b was calculated using the following formula:

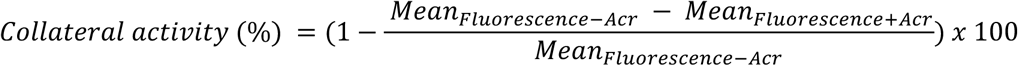

The standard deviation was calculated using the following formula:

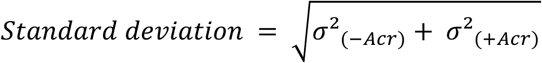

All custom code that was used for the analysis is available on GitHub (Github.com/Katwandu/AcrVIB1-characterization).

### Northern blotting analysis

For northern blotting analysis, 10 μg of each total RNA sample were mixed with 2x GLII loading buffer (NEB, Cat. no. B0363A), boiled at 95°C for 5 min in a closed vial and loaded on an 8% polyacrylamide gel containing 7 M urea for electrophoresis-based molecules separation at 300 V for 140 min using a gel transfer system (Doppel-Gelsystem Twin L, PerfectBlue-VWR).

After electrophoresis, RNA was transferred to Hybond-XL membranes (BLOTTING- NYLON 66 MEMBRANES,TYPE B, POS. 15356-1EA, Sigma-Aldrich) using an electroblotter with an applied voltage of 50 V for 1 h at 4°C (tank electroblotter Web M, PerfectBlue), cross- linked with UV light at 0.12 joules (UV lamp T8C; 254 nm, 8W), hybridized overnight in 15 mL of Roti Hybri-Quick buffer (A981.1, Carl Roth) at 42°C with 5 µL of γ ^32^P-ATP end-labeled oligodeoxyribonucleotides and visualized after one day of exposure on a phosphorimager (Typhoon FLA 7000, GE Healthcare). Oligonucleotide KW-pr-610 was used to generate the blot analysis in **Fig. 2C**. Loading uniformity was confirmed with the control-oligonucleotide binding to the ribosomal 5S RNA.

### Western blotting analysis

Samples for Western Blot were prepared by adding 1X Loading Dye and boiling them at 95°C for 10 min. Samples were loaded together with PageRuler Plus Prestained Protein Ladder (LIFE Technologies, #26619) in a 4-20% Mini-PROTEAN TGX Precast Protein Gel (Bio-Rad Laboratories, #4561095). The gels were placed in a clean plastic electrophoresis chamber and corresponding gel holder together with 1X SDS Running Buffer, the electrophoresis machine was connected to a power supply and all gels were run at 20 mA (60 V) for 20 min and then at 40 mA (120 V) for 45 min. Afterward, the gels were transferred to a Trans-Blot Turbo Mini 0.2 µm PVDF Transfer Pack (Bio-Rad Laboratories, #1704156) and blotted for 7 min using the Trans-Blot Turbo Transfer System. The blots were incubated in 5% nonfat dry milk prepared in 1X Tris-buffered saline with Tween 20 (TBS-T, VWR International, #J63682.K2) for 30 min at room temperature to block nonspecific binding sites. Primary antibodies were diluted in 5% nonfat dry milk according to the manufacturer’s protocol and added to the appropriate blots overnight at 4°C. All blots were washed three times for 20 min using 1X TBS-T before adding the appropriate secondary antibody. Blots were then incubated at room temperature for 1 h. Subsequently, blots were washed again three times for 5 min each using 1X TBS-T. The Odyssey CLx LI-COR was used for imaging.

### RNA labeling

For the 5′-end labeling of RNA with Cy5 used for EMSA, MST, RNA protection assay, synthetic oligoribonucleotides (IDT) were labeled with Cy5 maleimide conjugates (#GEPA15131, Sigma-Aldrich) using the 5′ EndTag kit (#VEC-MB-9001, BIOZOL) as directed by the manufacturer.

For the 3′-end labeling of RNA with Cy5 used for EMSA, crRNA processing assay, synthetic oligoribonucleotides (IDT) were treated with T4 RNA ligase (#EL0021, Thermo Fisher Scientific) in the presence of pCp-Cy5 (#NU-1706-CY5, Jena Bioscience). The labeled RNA was purified with RNA Clean and Concentrator columns-5 (#R1013, Zymo Research).

### Microscale thermophoresis (MST) and Spectral Shift

For the labeled AcrVIB1 binding experiments, AcrVIB1-His was labeled using the Monolith His-Tag Labeling Kit RED-tris-NTA 2nd Generation kit (Nanotemper, MO-L018) according to the manufacturer‘s instructions. For each binding experiment, AcrVIB1 was diluted to 20 nM using a binding buffer used previously for EMSA experiments. A series of 16 tubes with PbuCas13b or crRNA dilutions were prepared using the binding buffer, producing ligand concentrations ranging from 120 pM to 4 μM for PbuCas13b and 3 nM to 100 μM for crRNA. For measurements, each ligand dilution was mixed with one volume of labeled AcrVIB1, which led to a final concentration of 10 nM of labeled AcrVIB1 and 60 pM to 2 μM of Cas13b and 1.5 nM to 50 μM of crRNA. For the labeled RNA-binding experiments, labeled crRNA was diluted to 10 nM in the binding buffer. To form the RNP complex, 2 nM of Pbucas13b and 10 nM of labeled crRNA were pre-incubated for 15 min at room temperature in the binding buffer. To form the AcrVIB1:PbuCas13b complex, 105 μM of AcrVIB1 and 21 μM of PbuCas13b were pre-incubated for 15 min at room temperature in the binding buffer. A series of 16 tubes with PbuCas13b, AcrVIB1 or (AcrVIB1+PbuCas13b) complex dilutions were prepared in binding buffer, producing ligand concentrations ranging from 640 pM to 21 μM for PbuCas13b, 600 pM to 20 μM for AcrVIB1 and 300 pM to 10.5 μM for the complex. For the measurement, each PbuCas13b and complex ligand dilutions were mixed with one volume of labeled crRNA. Similarly, each AcrVIB1 dilution was mixed with one volume of the labeled RNP complex, which led to a final crRNA concentration of 5.0 nM in all reactions. When monitoring the RNA target binding, first the RNP complex was formed with 3 μM of unlabeled crRNA and 6 μM of PbuCas13b. The RNP was serially diluted and then 10 nM Cy5-5’-end-labeled target was added to each dilution yielding final concentrations of 22 pM - 1.5 μM of the RNP complex and 5 nM of labeled target RNA. Similar experiments were conducted to observe the effect of AcrVIB1 on target binding. Here, 6 μM PbuCas13b and 30 μM AcrVIB1 were pre-incubated for 15 min at room temperature in the binding buffer, followed by incubation with 3 μM of unlabeled crRNA for 15 min at room temperature.

Following the addition of the labeled targets to the ligand dilutions, the reactions were mixed by pipetting, incubated for 15 min at room temperature. In case of target RNA-binding, reactions were incubated for 15 min at 37°C. Capillary forces were used to load the samples into Monolith NT.115 Premium Capillaries (NanoTemper Technologies). Measurements were performed using a Monolith Pico instrument (NanoTemper Technologies) at an ambient temperature of 25°C. Instrument parameters were adjusted to 5% LED power, medium MST power, and MST on-time of 1.5 s. An initial fluorescence scan was performed across the capillaries to determine the sample quality and afterward, 16 subsequent thermophoresis measurements were performed. Data of three independently pipetted measurements were analyzed for the ΔF_norm_ values determined by the MO.Affinity Analysis software (NanoTemper Technologies). Graphs were plotted and binding affinities (linear regression model – one site specific model) were calculated using GraphPad Prism 9.2.0.

Binding affinities for AcrVIB1 titrated with the Helical-2 domain of PbuCas13b were obtained using a MonolithX instrument (NanoTemper Technologies) using both Spectral Shift and MST mode for evaluation. AcrVIB1 was fluorescently labeled using a Cy5-NHS ester by mixing 100 µL of a 20 µM protein solution with 100 µL of 60 µM dye. The reaction was carried out at room temperature for 30 minutes in 20 mM HEPES/NaOH pH 7.5, 150 mM NaCl. Excess dye was removed using a desalting column. The degree of labeling of AcrVIB1 with Cy5 was 30.8%. A dilution series of the non-labeled PbuCas13b Helical-2 domain was mixed with a constant concentration of Cy5-labeled AcrVIB1, resulting in 16 samples with 100 nM AcrVIB1 each and 50 µM Helical-2 domain as the highest ligand concentration in 20 mM HEPES/NaOH pH 7.5, 150 mM NaCl, 0.05% (v/v) Tween20. Affinity measurements were carried out in Premium Capillaries (NanoTemper Technologies) with 20% excitation power at 25°C. Binding was evaluated using both modes of the MonolithX device. In Spectral Shift mode, the 670 nm / 650 nm emission ratio of the initial fluorescence scans was evaluated to detect a bathochromic or hypsochromic shift of the dye induced by ligand binding. In MST mode, fluorescence was monitored at 670 nm and evaluated at 1.5 s on-time of the IR-laser at medium power. Data of four independent replicates were exported from the MO.Control2 software (NanoTemper Technologies) with conversion to ΔF_norm_ values for evaluation in MST mode. Graphs were plotted and binding affinities (nonlinear regression model – [Agonist] vs. response (three parameters)) were calculated using GraphPad Prism 10.3.0.

### Co-immunoprecipitation

To observe interactions between PbuCas13b-FLAG and AcrVIB1-His, Co-IP experiments were performed. For this, the purified labeled proteins were added together in 1x PBS at room temperature before adding them to Dynabeads Protein A labeled with either FLAG- or His- antibodies according to the manufacturer’s recommendations (LIFE Technologies, #10006D). First, the antibody for the target protein was incubated with the Dynabeads Protein A in a tube for 10 min. Excess antibody was washed away by placing the tube in a magnetic rack and removing the supernatant. To immunoprecipitate the target antigen, samples were mixed with the magnetic beads and rotated for 10 min at RT to allow the antigen to bind to the magnetic bead-Ab complex. The magnetic bead-Ab-Ag complex was washed three times using 200 μL of Washing Buffer for each wash step. After washing, the magnetic bead-Ab-Ag complex was resuspended in 100 μL of Washing Buffer and the bead suspension was transferred to a clean tube to avoid coelution of proteins bound to the tube wall. To elute the target antigen, 20 μL of Elution Buffer and 20 µL of 1X Protein Loading Buffer were added to the magnetic bead-Ab-Ag mix. After gently pipetting to resuspend the magnetic bead-Ab-Ag complex, the samples were boiled for 5 min at 95°C. The samples were subsequently used for western blotting analysis.

### crRNA processing assay

For the in vitro processing assay, 0.2 μM of PbuCas13b and 0.1 or 1 μM of AcrVIB1 were pre-incubated for 15 min at room temperature in a nuclease buffer (10 mM TrisHCl pH 7.5, 50 mM NaCl, 0.5 mM MgCl_2_, 0.1% BSA), then 5 nM Cy5-3′end-labeled extended crRNA (KW-RNA- 001) was added. The reaction was incubated for another 15 min at 37°C. All reactions were prepared in 10 μL and were stopped by adding 10 μL of colorless gel-loading solution (10 M urea, 1.5 mM EDTA, pH 8.0) on ice. The samples were separated on 10% PAAG-7M urea. The gels were visualized using an Amersham Typhoon imaging system. The assays were performed in two replicates.

### Cryo-electron microscopy Cryo-EM grid preparation

For the PbuCas13b:crRNA complex 1 mg/ml, for the PbuCas13b:AcrVIB1:crRNA complex 0.25 - 0.5 mg/ml and for the AcrVIB1:PbuCas13b Helical-2 domain complex 0.165 mg/ml SEC-purified samples were used for vitrification. 3.5-μL aliquots of each sample were applied to holey carbon grids (Quantifoil Cu R1.2/1.3, 200 mesh) that were glow discharged for 100 s at a current of 15 mA using a PELCO easiGlow^TM^ glow discharger (Ted Pella, Inc). The grids were then blotted once with filter paper for 4.5 sec to remove excess liquid and plunge-frozen in liquid ethane using a FEI Vitrobot Mark IV (Thermo Fisher Scientific Ltd) at 4 - 10°C and 100% humidity.

### Cryo-EM Data Acquisition

All cryo-EM data presented here were collected on a Glacios TEM equipped with a Selectris energy filter, and data collection parameters are detailed in Table S1. The microscope was operated in EFTEM mode with a slit-width of 10 eV and using a 70 μm objective aperture. Automated data acquisition was performed using EPU (Thermo Fisher Scientific). For the PbuCas13b:crRNA complex, 3,778 movies were collected, and for the PbuCas13b:AcrVIB1:crRNA complex, several datasets totaling 10,707 movies were collected. 8206 movies were collected for the AcrVIB1:PbuCas13b Helical-2 domain complex.

### Image Processing

Cryo-EM data analysis was conducted using cryoSPARC (version 4.5.3)^52–54^. For the PbuCas13b:crRNA complex, standard procedures including CTF correction, motion correction, and particle picking were employed. Over 3.3 million particles were picked by the template picker and initially classified by a hetero refinement. The best class, comprising around 1.9 million particles, underwent further 2D classification to remove debris and broken particles. From the remaining 1 million particles, iterative particle cleaning was achieved through multiple rounds of heterogeneous refinement and 2D classifications. For the final set of 356,629 cleaned particles, reference-based motion correction was performed followed by non-uniform refinement in cryoSPARC. The final reconstruction achieved an overall resolution of 3.09 Å, determined by Fourier shell correlation at a 0.143 cut-off.

For the PbuCas13b:AcrVIB1:crRNA datasets, the same processing route was followed. In template-based particle picking, over 5.4 million particles were extracted. These underwent classification in a heterogeneous refinement, with the largest class containing 1.25 million particles. Subsequent 2D classification revealed a mixture of closed and open particle views. The subclass of the open form of PbuCas13b:AcrVIB1:crRNA was separated from the closed form through several rounds of 2D classifications and a final 3D classification. The remaining 302,661 particles were further classified by heterogeneous refinements and 3D classifications into three final subclasses of the open form of PbuCas13b:AcrVIB1:crRNA. The particle numbers for the final subclasses ranged from 74,499 to 120,153 particles per class. The final reconstructions of each subclass had an overall resolution ranging from 4.23 Å to 4.49 Å, determined by Fourier shell correlation at a 0.143 cut-off.

The initial AcrVIB1:PbuCas13b Helical-2 domain complex dataset comprised approx. 7.2 Mio particles. With a subset of this data, six ab-initio models were created and used to classify the 7.2 Mio particles by a heterogeneous refinement followed by a non-uniform refinement. The best model/particles were subjected to several iterative 2D classifications and non-uniform refinements to result in a 1,078,020 large particle set that refined to 5.11 Å as determined by Fourier shell correlation at a 0.143 cut-off. Cryo-EM data and refinement statistics are summarized in **Table S1**.

### Structure refinement and model building

The crystal structure model of a PbuCas13b:crRNA complex (PDB: 6DTD) served as initial template and was rigid-body fitted into the cryo-EM density within UCSF ChimeraX^55,56^ using Isolde^57^, and manually adjusted and rebuilt in Coot^58^. Several rounds of real-space refinement were performed in the PHENIX software suite^59,60^ before final validation. The final structure was refined and validated before being deposited into the protein database with the PDB code 9FCV. The final electron density map was deposited at EMDB with the accession code EMD- 50322.

For the various flexible open forms of the PbuCas13b:AcrVIB1:crRNA complex, the final model of the PbuCas13b:crRNA complex was superposed onto the resulting maps of the three subclasses. The flexible arm region and the crRNA 5′ end were manually adjusted and fitted with Isolde due to low and poor density in this region. No real-space refinement was performed, and no final model of the corresponding subclasses was deposited, but the final electron density map with the accession code EMD-51509.

The AlphaFold3 derived model of the AcrVIB1:PbuCas13b Helical-2 domain complex was superposed into the map of the AcrVIB1:PbuCas13b Helical-2 domain complex using UCSF ChimeraX “fit in map” base tool functionality without further model modifications applied. The final map was deposited and has the accession code EMD-51513.

### Electrophoretic mobility shift assays

For the electrophoretic mobility shift assays (EMSAs), first 0.2 μM PbuCas13b and 0.1 or 1 μM AcrVIB1 were pre-incubated for 15 min at room temperature in a binding buffer (10 mM TrisHCl pH 7.5, 50 mM NaCl, 0.1% BSA, 10 mM EDTA, 5% glycerol) in the presence of 1 μg/μL of yeast RNA (#AM2283, Ambion) to enhance specific RNA-protein interaction, then 5 nM of the Cy5-5′-labeled crRNA (KW-RNA-002) was added. The reaction was incubated for another 15 min at room temperature.

When monitoring the target RNA binding, first 100 nM unlabeled crRNA (KW-RNA- 002) was incubated for 15 min at room temperature with the pre-incubated proteins, then 5 nM Cy5-3′-labeled target (KW-RNA-005) was added. The reaction was incubated for another 15 min at 37°C.

The samples were separated on a 2% agarose-TBE gel at 4°C. The gels were visualized using an Amersham Typhoon imaging system. The assays were performed in two replicates.

### RNA protection assays

For the RNA protection assay, 1 μM of PbuCas13b and 5 μM of AcrVIB1 were pre-incubated for 15 min at room temperature in a cleavage buffer (10 mM HEPES pH 7.0, 250 mM NaCl, 5 mM MgCl_2_, 1 mM DTT, 0.05% Tween 20) in the presence of 1 μg/μL of yeast RNA (#AM2283, Ambion), then 5 nM of the Cy5-5′-labeled crRNA (KW-RNA-002) was added. After the reaction was incubated for another 15 min at room temperature, it was treated with the 1 U/μL of RNase T1 (#AM2283, Ambion) for 30 min at 37°C. All reactions were prepared in a total volume of 10 μL and were stopped by adding 10 μL of colorless gel-loading solution (10 M urea, 1.5 mM EDTA, pH 8.0) on ice. The samples were separated on a 6% PAA-7M urea gel. The gels were visualized using an Amersham Typhoon imaging system. The assays were performed in two replicates. The data was analyzed with GraphPad Prism 9.3.0.

### Complex generation, stability and size analyses

Complexes of full-length PbuCas13b were formed in 100-µl samples by incubation of the protein (5 µM) with 1.5-fold molar excess of ligands. For complexes containing either unprocessed cRNA (KW-RNA-001) or AcrVIB1, the samples were incubated for 1 hour at room temperature after addition of the corresponding ligand. For complexes with three components, a second ligand (either AcrVIB1 or cRNA) was added for overnight incubation at 4°C. Samples were after incubation centrifuged for 10 min at maximum speed in a benchtop microcentrifuge.

The complex of AcrVIB1 and the Helical-2 domain of PbuCas13b was formed by mixing equimolar amounts of both proteins and subjecting the mixture directly onto a Superdex 75 gel filtration column of appropriate volume equilibrated in 20 mM HEPES/NaOH pH 7.5, 150 mM NaCl.

Stability analyses were performed in a Prometheus Panta instrument (Nanotemper). Samples were in triplicates heated up to 95°C from 25°C at 3°C/min. Unfolding/Aggregation was followed by observing changes in the intrinsic fluorescence of aromatic residues up on unfolding of the proteins (350/330 nm fluorescence ratio), turbidity measured by static light scattering and the increase in cumulant hydrodynamic radius by dynamic light scattering.

Sizes of the full-length PbuCas13b complexes were determined by mass photometry in a TwoMP instrument (Refeyn Ltd). The samples were diluted to 100 nM and 1 µL of sample was added to 9 µL of buffer on the glass slide of the mass photometer. Movies of 90-s length were recorded for each sample at normal field of view in AcquireMP (Refeyn Ltd). Evaluation was performed in DiscoverMP (Refeyn Ltd) on the last 1500 counts recorded in each movie. Molecular masses were determined by matching histogram contrast against a molecular weight standard (Invitrogen/Thermo NativeMarkTM, LC0725).

### SEC-MALS analysis

For analysis of the total mass of AcrVIB1 in complex with the Helical-2 domain of PbuCas13b we applied analytical size exclusion chromatography in combination with multi angle light scattering (SEC-MALS). Therefore, we applied 135 ug protein to a Superdex 75 Increase 10/300 GL column (Cytiva) coupled to our MALS-System, comprising an Agilent 1260 Infinity II HPLC system (Santa Clara, USA), a mini-DAWN TREOS II multi-angle laser light-scattering detector (Wyatt, Santa Barbara, USA) and an Optilab dRI detector (Wyatt, Santa Barbara, USA). The column was pre-equilibrated in 20 mM HEPES pH 7.5 and 150 mM NaCl. Data analysis was carried out with Astra Version 8.2.2. For molecular mass calculation a dn/dc of 0.185 mL/g was assumed.

### TXTL assays

Plasmids encoding either the nuclease, a crRNA, deGFP or AcrVIB1 were used to assess the targeting activity of the nuclease and the inhibitory activity of AcrVIB1 in a cell-free transcription-translation (TXTL) assay. The nuclease was under the control of a T7 promoter; therefore a plasmid encoding the T7 RNA polymerase had to be added. The crRNA encoded either a targeting (T) spacer or non-targeting (NT) control. The assay is described in full previously^25^. In brief, we pre-expressed the AcrVIB1 plasmid in TXTL for 16 h with a final concentration of 4 nM, before adding the pre-expressed protein to fresh TXTL mix together with the plasmids encoding the Cas13b nuclease, a crRNA, T7RNAP or deGFP with a final concentration of 1 nM for the nuclease, the crRNA and deGFP and 0.2 nM T7RNAP. The samples were then incubated at 29°C for 16 h in a plate reader (BioTek Synergy H1), and fluorescence was measured every three minutes (excitation, emission: 485 nm, 528 nm). All TXTL reactions were prepared using the Echo525 Liquid Handling system to distribute the plasmids. The TXTL mix was distributed manually beforehand. Herein, a set of plasmids containing single amino-acid substitutions in either PbuCas13b or AcrVIB1 were tested for the impact on targeting and inhibitory activity, respectively. Mutations of PbuCas13b included K627A, N634A, and R662A. AcrVIB1 was mutated at three different sites, D37/38A, S56A, T70A. Furthermore, we tested four amino-acid substitutions in RanCas13b to test if they confer sensitivity to AcrVIB1 to the previously unaffected RanCas13b nuclease, K559N, Y609K, W627R and R630Q.

### Statistical analyses

In this study, unpaired, one-sided t-tests, with the incorporation of Welch’s correction, were employed to evaluate the statistical significance of observed differences between two independent groups. We employed these analyses for obtaining the statistical significance for crRNA degradation by RNase T1 and for binding measurements via MST.

## Supplementary Figures

**Figure S1.**
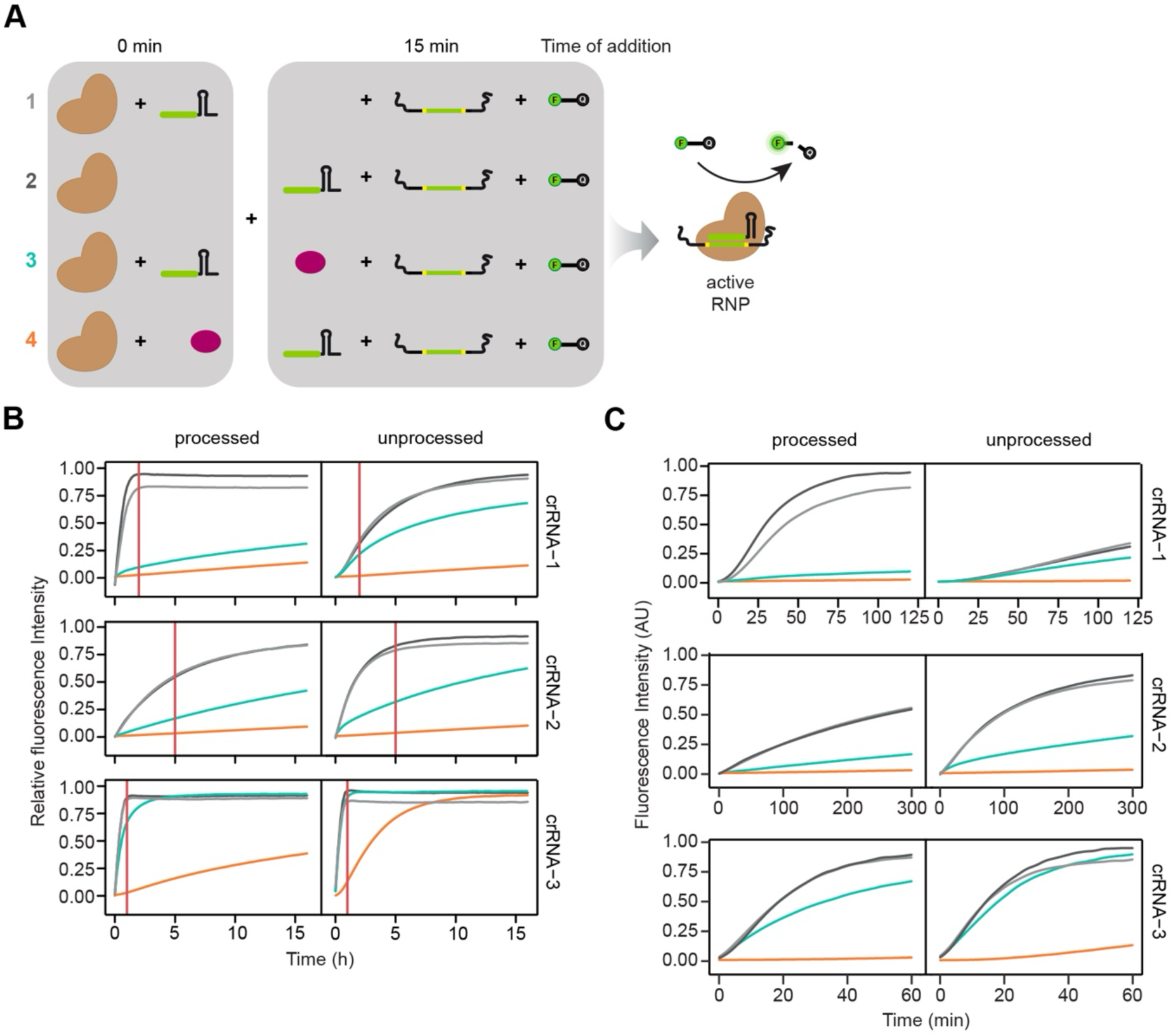
AcrVIB1 inhibits PbuCas13b more efficiently prior to RNP complex formation. (**A**) PbuCas13b was incubated with either a corresponding crRNA or AcrVIB1 for 15 minutes, before all other components were added. A fluorescently labeled FAM reporter and a target RNA matching the crRNA were always added last. Fluorescence was measured over time. (**B**) Full time courses of collateral RNA cleavage assays. The red line indicates the endpoint chosen for each crRNA. See Fig. 1 for corresponding data. (**C**) Time courses showing fluorescence measurements using endpoints of 2 h, 5 h and 1 h for crRNA-1, 2 and 3, respectively.

**Figure S2.**
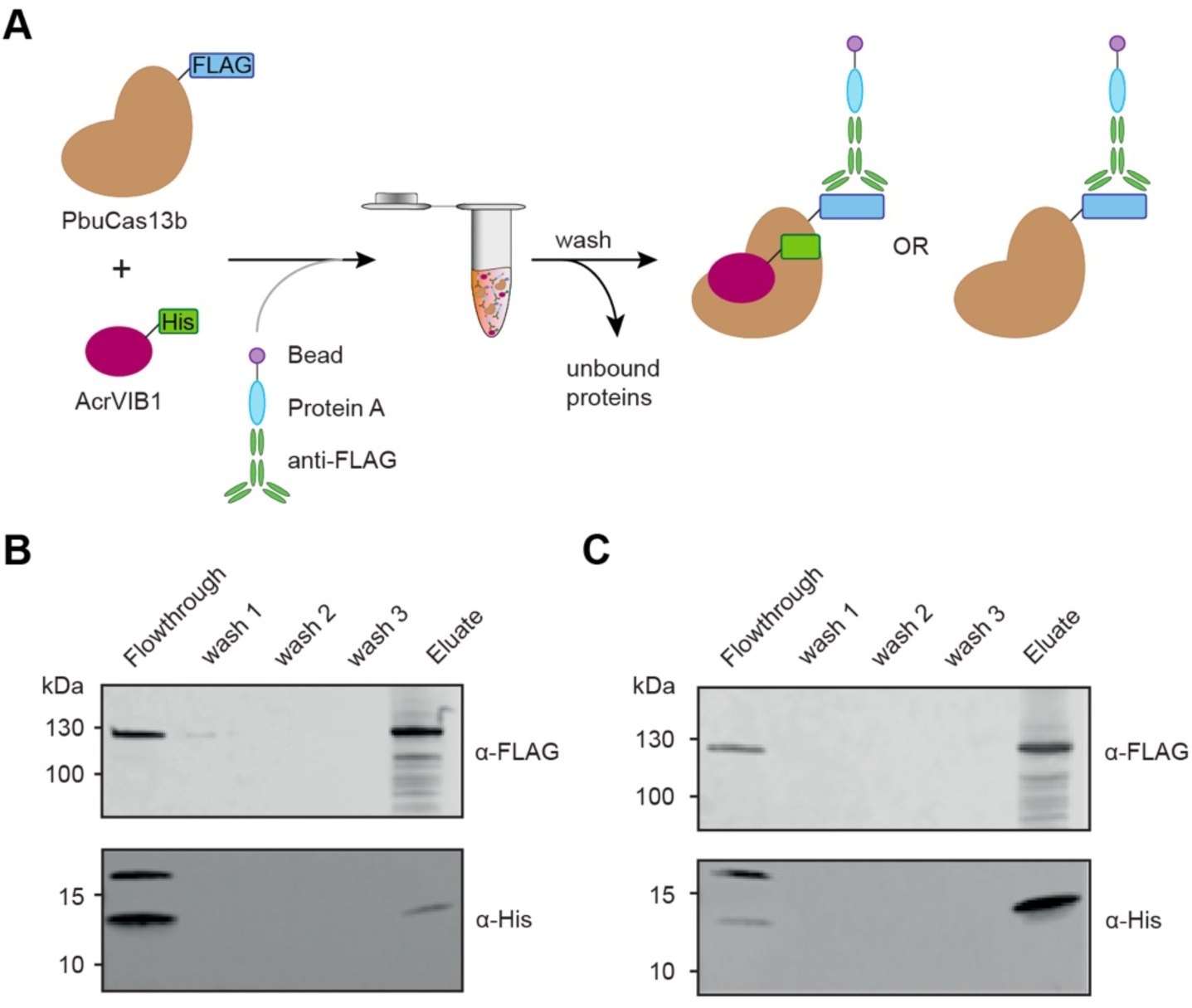
AcrVIB1 and PbuCas13b directly interact. Co-immunoprecipitation shows interaction between AcrVIB1 and PbuCas13b. Incubation of both proteins with subsequent treatment with antibody-labeled magnetic beads leads to co-elution of both proteins in the elution fraction after three washing steps. Gel images in B and C are representative of two replicates.

**Figure S3.**
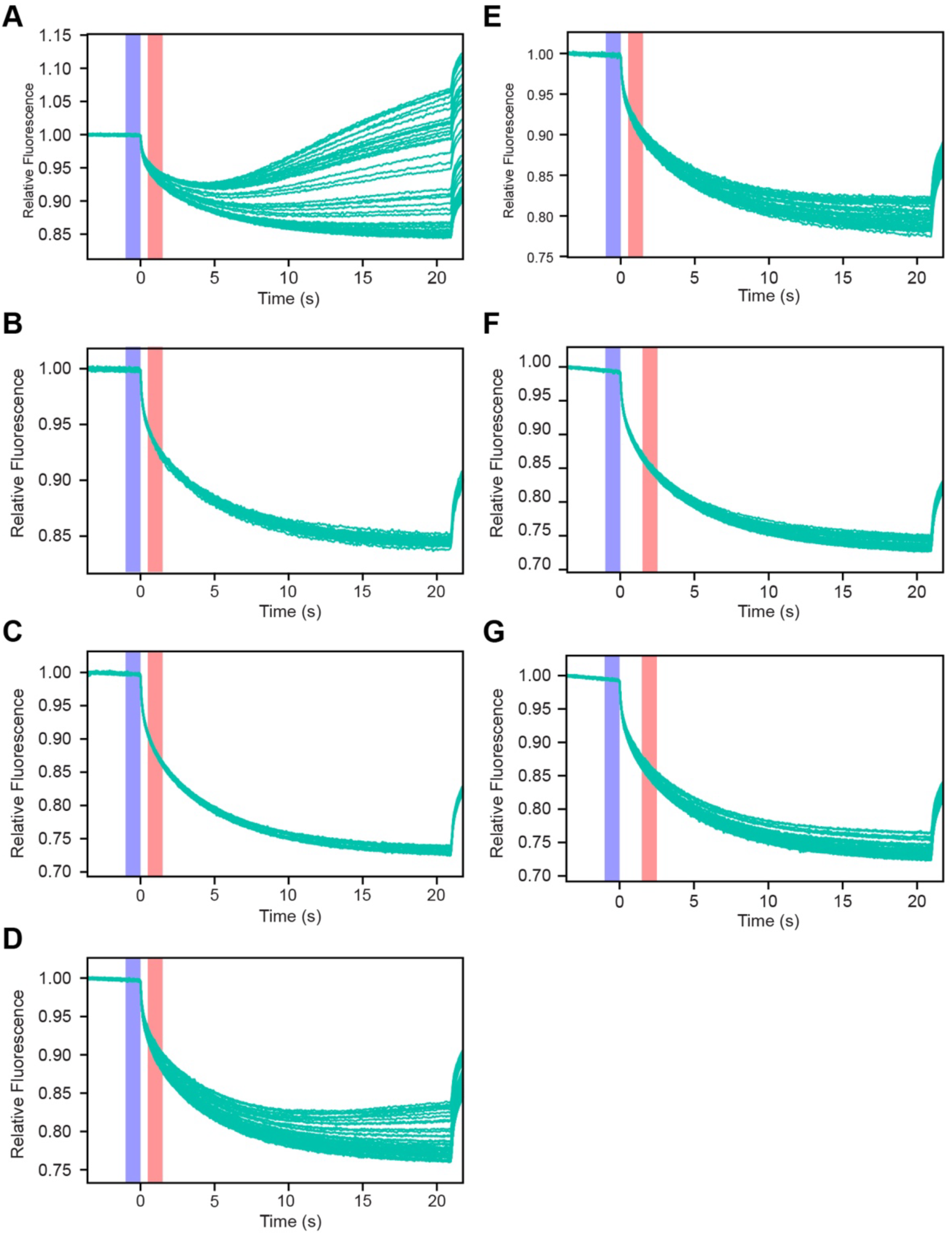
Thermophoretic time traces of microscale thermophoresis (MST) measurements. Each panel captures binding between two designated components. (**A**) PbuCas13b and labeled AcrVIB1. (**B**) crRNA and labeled AcrVIB1. (**C**) AcrVlB1 and labeled RNP. (**D**) PbuCas13b and labeled crRNA. (**E**) PbuCas13b:AcrVIB1 complex and labeled crRNA. (**F**) RNP and labeled target RNA. (**G**) (PbuCas13b:AcrVIB1:crRNA) complex and labeled target RNA. Blue and pink boxes in the time-course traces represent the temperature jump and MST-on time (1.5 s), respectively. In all cases, there was no adsorption of the labeled protein to the capillaries and no protein aggregation was observed. See Figures 2 and 3 for resulting binding curves.

**Figure S4.**
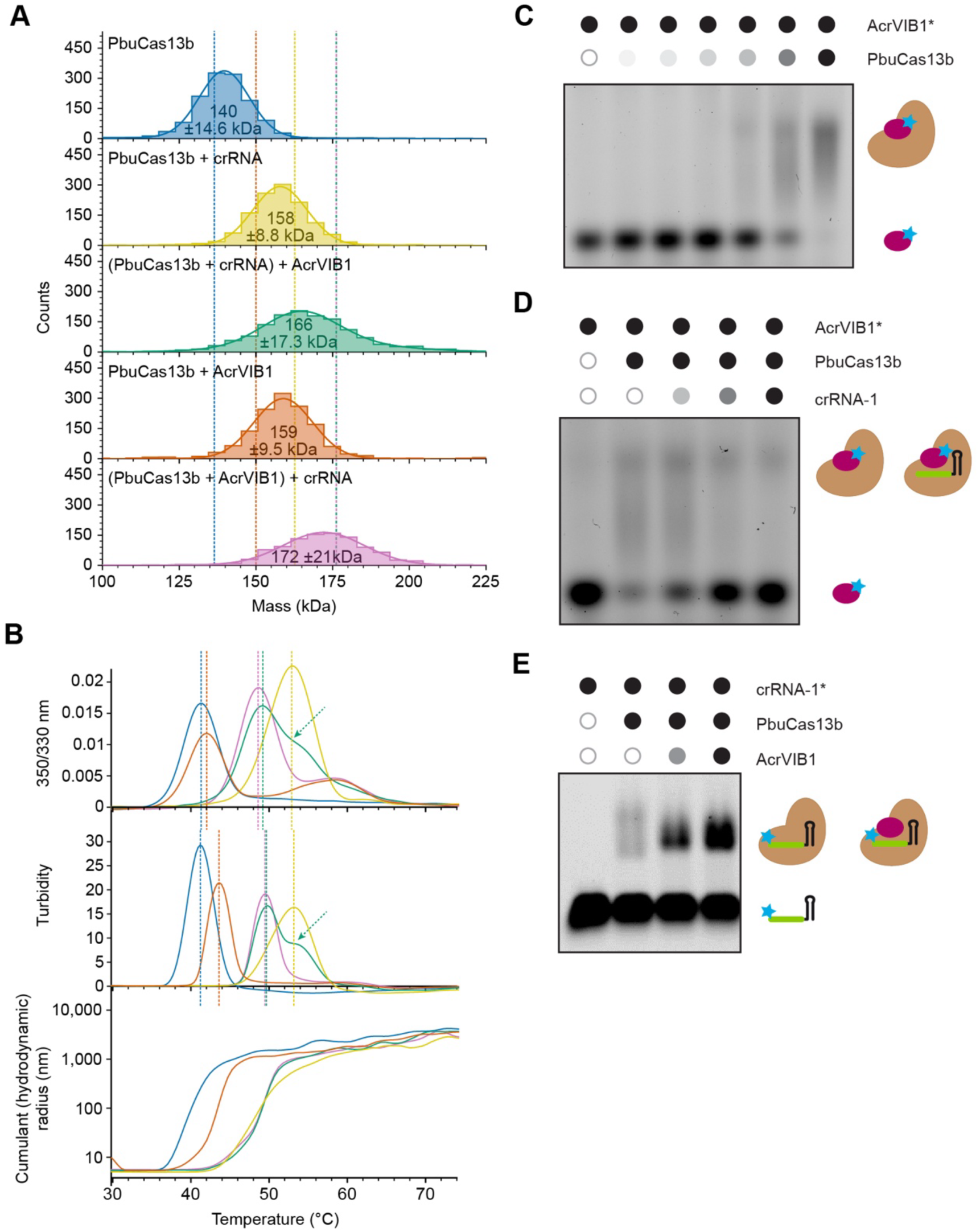
AcrVIB1 principally binds PbuCas13b, with binding promoting crRNA binding. (**A**) Mass photometry analysis of PbuCas13b in complex with AcrVIB1 and/or a 81-nt crRNA. The indicated molecular masses were determined by Gaussian fits to the histograms. Dashed lines indicate theoretical masses of the corresponding complexes. Blue: PbuCas13 only, yellow: PbuCas13b + RNA (1 h RT), green: PbuCas13b + RNA (1h RT) + AcrVIB1 (4°C o/n), orange: PbuCas13b + AcrVIB1 (1 h RT), violet: PbuCas13b + AcrVIB1 (1h RT) + RNA (4°C o/n). (**B**) Melting curve determination of the same samples as in (A). Unfolding/aggregation was followed by observing changes in the intrinsic fluorescence of aromatic residues (ratio of 350/330 nm, first derivative, top), turbidity measurement (first derivative, middle) and dynamic light scattering (cumulant hydrodynamic radius, bottom). Melting temperatures derived from fluorescence and turbidity measurements are shown as dashed lines. Blue: PbuCas13 only, yellow: PbuCas13b + RNA (1h RT), green: PbuCas13b + RNA (1 h RT) + AcrVIB1 (4°C o/n), orange: PbuCas13b + AcrVIB1 (1 h RT), violet: PbuCas13b + AcrVIB1 (1 h RT) + RNA (4°C o/n). The green arrow indicates a shoulder of the PbuCas13b + RNA + AcrVIB1 sample that reflects a significant fraction of PbuCas13b + RNA without bound AcrVIB1. Plots show the mean of 3 replicates. (**C**) EMSA evaluating AcrVIB1 binding to PbuCas13b. AcrVIB1 (5 nM) was labeled through its fused C-terminal His-tag. PbuCas13b was added in six concentrations, left to right from lowest to highest (5 nM, 10 nM, 50 nM, 100 nM, 200 nM, 1,000 nM). (**D**) EMSA examining the influence of crRNA on AcrVIB1 binding to PbuCas13b. AcrVIB1 (5 nM) was labeled at the His-tag for visualization on the gel. crRNA-1 was incubated with PbuCas13b (200 nM) before the addition of AcrVIB1. Three concentrations of crRNA-1 were used: 0.1 µM, 1 µM and 10 µM. (**E**) EMSA assessing 5′-labeled crRNA (5 nM) binding to PbuCas13b (200 nM) with and without AcrVIB1. AcrVIB1 was added in two concentrations (100 nM and 1 µM, indicated by the gray and black circle, respectively). Gels shown in C, D & E are representative of two independent replicates.

**Figure S5.**
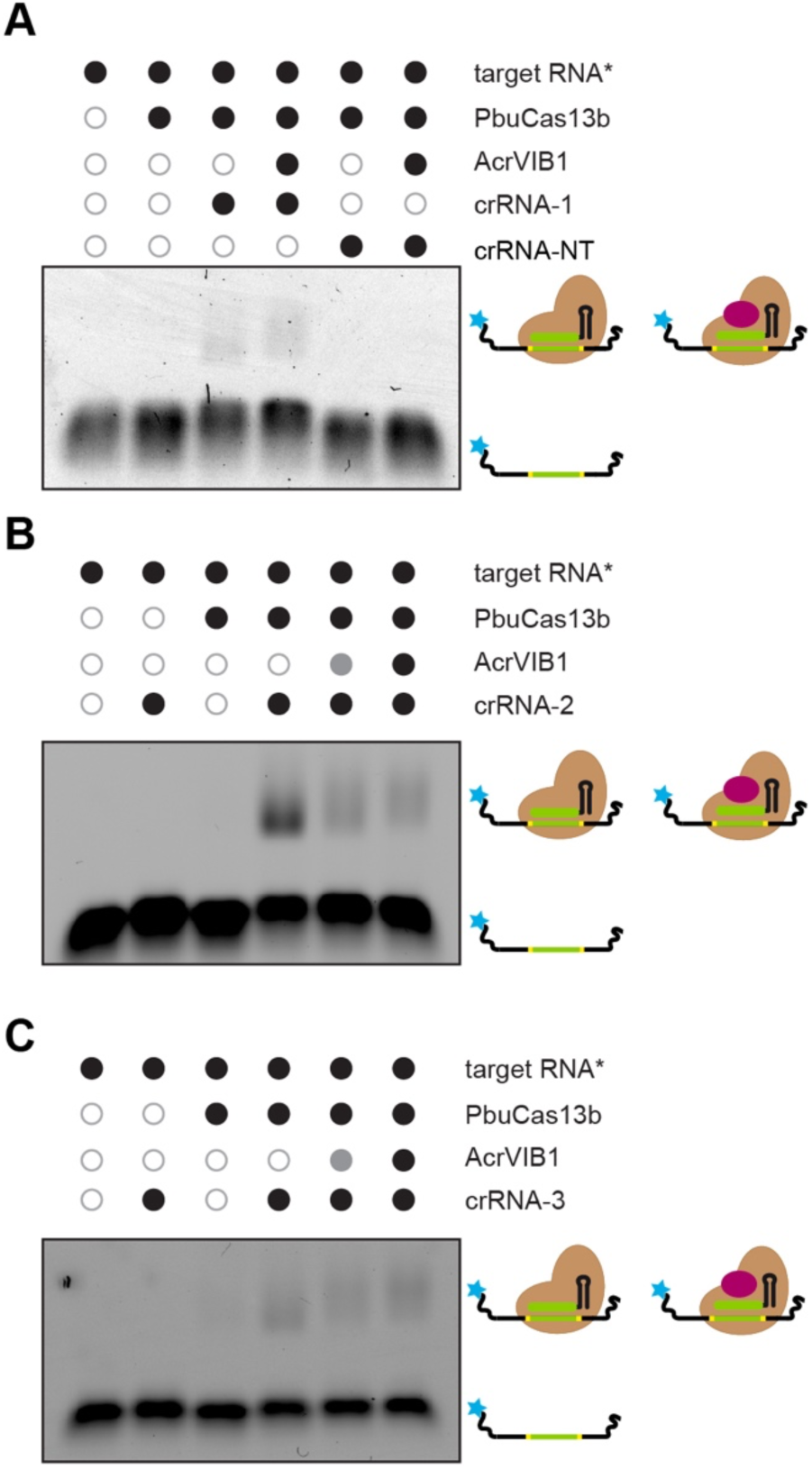
AcrVIB1 alters target binding to the PbuCas13b:crRNA complex. (**A**) EMSA assessing 5′-labeled target RNA (5 nM) binding to PbuCas13b:crRNA-1 (200 nM and 100 nM, respectively) with and without AcrVIB1 (1,000 nM). (**B**) EMSA assessing 5′-labeled target RNA (5 nM) binding to PbuCas13b:crRNA-2 with or without AcrVIB1. (**C**) EMSA assessing 5′-labeled target RNA (5 nM) binding to PbuCas13b:crRNA-3 with or without AcrVIB1. In B and C, AcrVIB1 was added at two final concentrations: 100 nM (gray circle) and 1,000 nM (black circle). RNP complexes were formed using 200 nM of PbuCa13b and 100 nM of crRNA. Gels shown in A-C are representative of two independent replicates.

**Figure S6.**
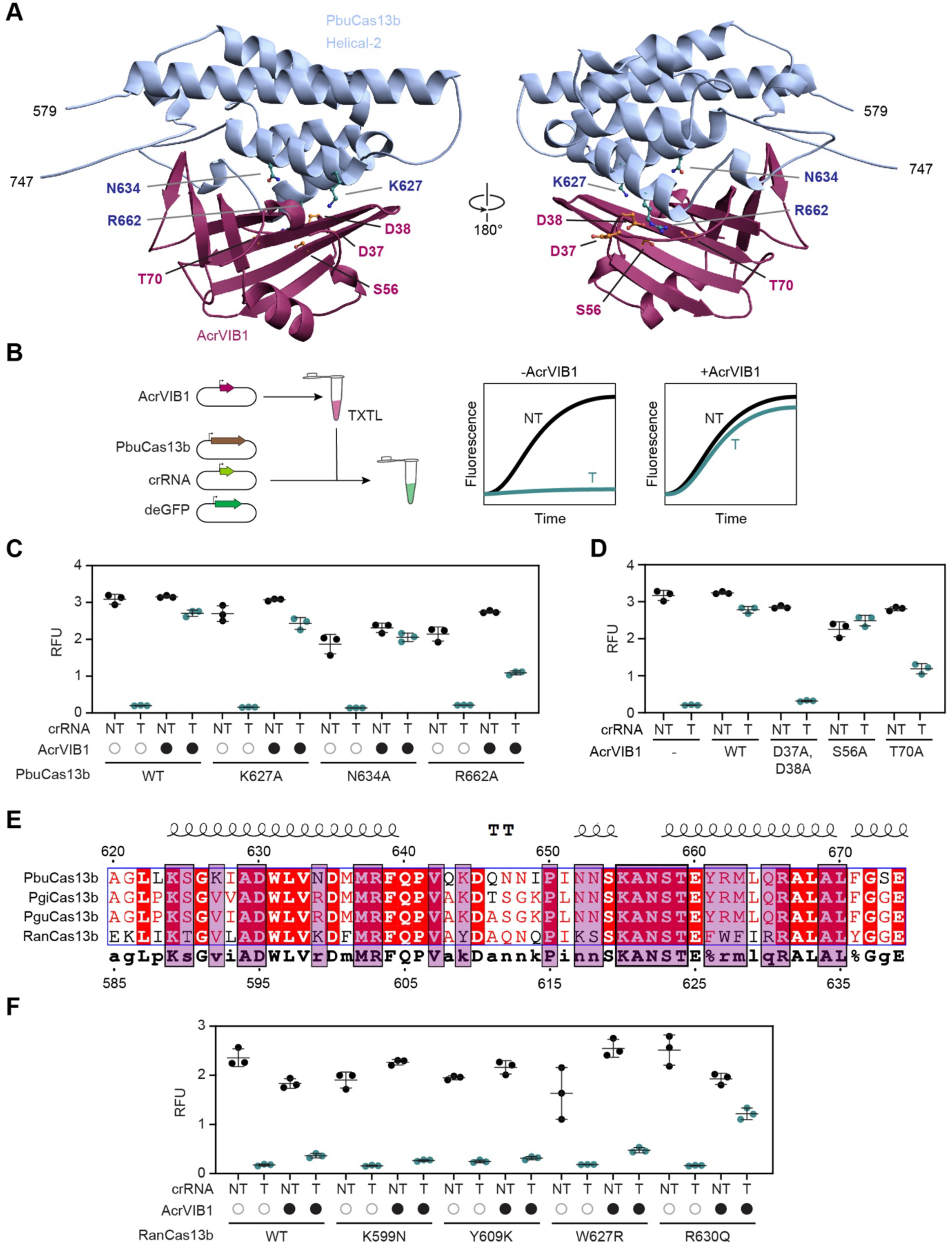
Impact of selected PbuCas13b, AcrVIB1 and RanCas13b mutations on inhibition in a cell-free transcription-translation (TXTL) assay. (**A**) Model of the binding interface between AcrVIB1 and the Helical-2 domain of PbuCas13b shown in Figure 6B. Selected residues in PbuCas13b and AcrVIB1 for mutational analysis are shown. (**B**) Depiction of the TXTL assay. AcrVIB1 pre-expressed for 16 hours is combined with DNA constructs encoding PbuCas13b, a targeting (T) or non-targeting (NT) gRNA, and a targeted deGFP expression construct. (**C**) Impact of mutations to PbuCas13b in the TXTL assay. (**D**) Impact of mutations to AcrVIB1 in the TXTL assay. (**E**) Alignment of the Helical-2 domain across related Cas13b orthologs. (**F**) Impact of mutations to RanCas13b to match the corresponding residue in PbuCas13b in the TXTL assay. For C, D, and F, end-point measurements after 16 hours are shown. Circles represent individual measurements. Horizontal bars and error bars represent the mean and S.D. of three independent measurements.

**Table S1.** Cryo-EM data collection, structure refinement, and validation statistics.

| **Data collection and processing** | ***PbuCas13b:***<br>***crRNA complex***<br>**(PDB: 9FCV**<br>**EMDB: EMD-50322)** | ***AcrVIB1:PbuCas13b:***<br>***crRNA complex***<br>**(EMDB: EMD-51509)** | ***AcrVIB1:***<br>***PbuCas13b Helical-2***<br>***domain* (EMDB:**<br>**EMD-51513)** |
| --- | --- | --- | --- |
| Magnification | 130000 | 130000 | 130000 |
| Voltage (kV) | 200 | 200 | 200 |
| Electron exposure (e-/Å<sup>2</sup>) | 40 | 40 | 60 |
| Defocus range (µm) | -0.6 to -2.0 | -0.6 to -2.0 | -0.6 to -1.6 |
| Pixel size (Å) | 0.91 | 0.91 | 0.91 |
| Symmetry imposed | C1 | C1 | C1 |
| Image stacks (no.) | 3778 | 10707 | 8206 |
| Initial particle images (no.) | 1903944 | 1250433 | 7229692 |
| Final particle images (no.) | 356629 | 120153 | 1078020 |
| Map resolution (Å) | 3.09 | 4.23 | 5.11 |
| FSC threshold | 0.143 | 0.143 | 0.143 |
| **Refinement** |  |  |  |
| No. of protein residues | 1057 |  |  |
| No. of RNA bases | 44 |  |  |
| No. of atoms | 19225 |  |  |
| **RMSDs** |  |  |  |
| Bond lengths (Å) | 0.002 |  |  |
| Bond angles (°) | 0.404 |  |  |
| **PDB validation** |  |  |  |
| Clash score | 2.34 |  |  |
| Poor rotamers (%) | 0.00 |  |  |
| C-Beta deviation | 0.00 |  |  |
| **Ramachandran plot** |  |  |  |
| Favored (%) | 97.7 |  |  |
| Allowed (%) | 2.2 |  |  |
| Disallowed (%) | 0.1 |  |  |

